# Medial entorhinal cortex plays a specialized role in learning of flexible, context-dependent interval timing behavior

**DOI:** 10.1101/2023.01.18.524598

**Authors:** Erin R. Bigus, Hyun-Woo Lee, John C. Bowler, Jiani Shi, James G. Heys

## Abstract

Episodic memory requires encoding the temporal structure of experience and relies on brain circuits in the medial temporal lobe, including the medial entorhinal cortex (MEC). Recent studies have identified MEC ’time cells’, which fire at specific moments during interval timing tasks, collectively tiling the entire timing period. It has been hypothesized that MEC time cells could provide temporal information necessary for episodic memories, yet it remains unknown whether MEC time cells display learning dynamics required for encoding different temporal contexts. To explore this, we developed a novel behavioral paradigm that requires distinguishing temporal contexts. Combined with methods for cellular resolution calcium imaging, we find that MEC time cells display context-dependent neural activity that emerges with task learning. Through chemogenetic inactivation we find that MEC activity is necessary for learning of context-dependent interval timing behavior. Finally, we find evidence of a common circuit mechanism that could drive sequential activity of both time cells and spatially selective neurons in MEC. Our work suggests that the clock-like firing of MEC time cells can be modulated by learning, allowing the tracking of various temporal structures that emerge through experience.

## Main Text

Our daily experiences unfold across space and time, meaning the brain must capture these dimensions to accurately form episodic memories (i.e. memories of personal experiences that occur in a specific spatial and temporal context)^1,2^. Medial temporal lobe (MTL) structures are critical for episodic memory, raising the question of how these regions encode space and time. A remarkable series of findings have revealed the role of MTL structures in encoding space, beginning with the findings that 1) MTL regions are critical in memory-guided spatial navigation behavior^3–6^ and 2) contain so-called place cells in the hippocampus^5,7^ and grid cells in the medial entorhinal cortex (MEC)^8^ that fire when animals visit particular locations with an environment. Critically, 3) spatial cells remap, reorganizing their firing fields as animals navigate and learn features of different environments to form a unique map for each spatial context^9,10^. Observing such learning dynamics provided the invaluable insight that a key role of spatially tuned cells is likely to create a “cognitive map” of an environment^11,12^ that can be stored in memory and used to guide future behavior.

In contrast to spatial context, it remains relatively unknown how temporal context, or the temporal structure of experiences, is encoded within the MTL memory system. The nervous system must track time across many scales, ranging from milliseconds to hours, but the intermediate scale of interval timing (seconds to minutes) is perhaps most relevant for planning and executing daily behaviors including foraging, mating, prey capture and avoidance^13–15^. Accordingly, encoding the temporal structure of daily experiences requires interval timing. Though it’s largely unclear how the passage of interval time and the duration of events are tracked and recorded within the MTL, one intriguing possibility is that common mechanisms support encoding of both space and time^16^. If so, as for space, we would expect MTL regions to 1) be necessary for interval timing behavior, 2) contain cells selective to time, and 3) use distinct patterns of time-selective cells to form “timelines” of unique experiences, akin to maps. Prior work suggests that MEC fits the first two criteria: MEC is both 1) necessary for interval timing behavior^17–19^ and 2) contains time cells that fire regularly at discrete moments as rodents report temporal durations on the scale of seconds^20^. As a population, different MEC time cells fire regularly at different moments in a timed interval, like the second hand of a clock, thereby forming a sequence of neural activity, tiling the entire timing epoch. By analogy to spatial cells, MEC time cells could play a key role in episodic memory by 3) using unique time cell trajectories to form distinct “maps”, or “timelines” of temporal experiences (i.e. contexts). This third point makes clear predictions about the learning dynamics of MEC time cells: distinct patterns of time cells should emerge as animals learn the temporal structure of an experience, and emergence of these patterns should be necessary for timing behavior. Evidence of such learning dynamics could suggest that MEC time cells play a key role in the formation of episodic memories by encoding the temporal structure of experiences. However, these predictions have yet to be experimentally tested.

We therefore aimed to test the hypotheses that 1) distinct populations of time cells will become active as animals learn to identify a new temporal context, forming a unique map or “timeline” of each temporal context, and 2) such dynamics support learning of timing behavior. To address these questions, we developed a novel temporal delayed nonmatch to sample (tDNMS) task that requires mice to differentiate the temporal structure of trials (temporal context). By performing 2-photon calcium imaging as mice performed the tDNMS task, we uncovered populations of MEC time cells that fire selectively at specific moments in the timing task, with the population of time cells creating a sequence that spans the entire timing epoch. Remarkably, we find that over the course of learning, these sequences become context-dependent, whereby specific populations of MEC time cells become active on particular trial types. Further, multiple lines of evidence suggest that the activity of MEC time cells plays a causal role specifically in learning context-dependent interval timing behavior. Finally, we find evidence for a common circuit mechanism that may support both MEC spatial and time coding. Our results suggest that MEC time cells may play a central role in episodic memory by forming unique “timelines” that encode the temporal structure of distinct experiences.

## Results

### Mice learn novel tDNMS task using flexible timing behavior

We designed a timing task with two objectives. First, mice must track time and make decisions based upon the temporal structure of each trial. Additionally, the task should require cognitive flexibility to maximally engage the MTL memory system. This second point may be critical to elicit learning dynamics, given prior work demonstrating that learning flexible but not rigid navigation behavior requires the MTL^21^. To meet these two goals, we adapted the delayed nonmatch to sample (DNMS) task structure, known to engage the MTL^22–24^, to create a novel temporal delayed nonmatch to sample (tDNMS) task. Because mice heavily rely on olfaction, we built a flow dilution olfactometer (Fig. S1a)^25^ and signaled stimuli via a single odorant (isoamyl acetate). We validated our system by ensuring that the concentration of the odorant remains constant over the course of 45 mins (a full training session) and that odor concentration can be rapidly controlled (Fig. S1b).

In each trial of the tDNMS task, a water-restricted, head-fixed mouse (Fig. 1a) is presented with two successive stimuli for either a short (2s) or long (5s) duration, separated by a brief 3s interstimulus interval (ISI). Trials are performed in the dark and are separated using 1) a 16-24s intertrial interval (ITI) and 2) a light pulse (0.25s) to signal trial start (Fig. S1c). The tDNMS task consists of three trial types defined by each stimulus duration: short-long (S-L), long-short (L-S), and short-short (S-S). Using a “Go/No-Go” strategy, mice are trained to lick to report a nonmatch of durations (Go trials; S-L & L-S) and withhold from licking in response to match durations (No-Go trials; S-S) (Fig. 1c).

**Figure 1.**
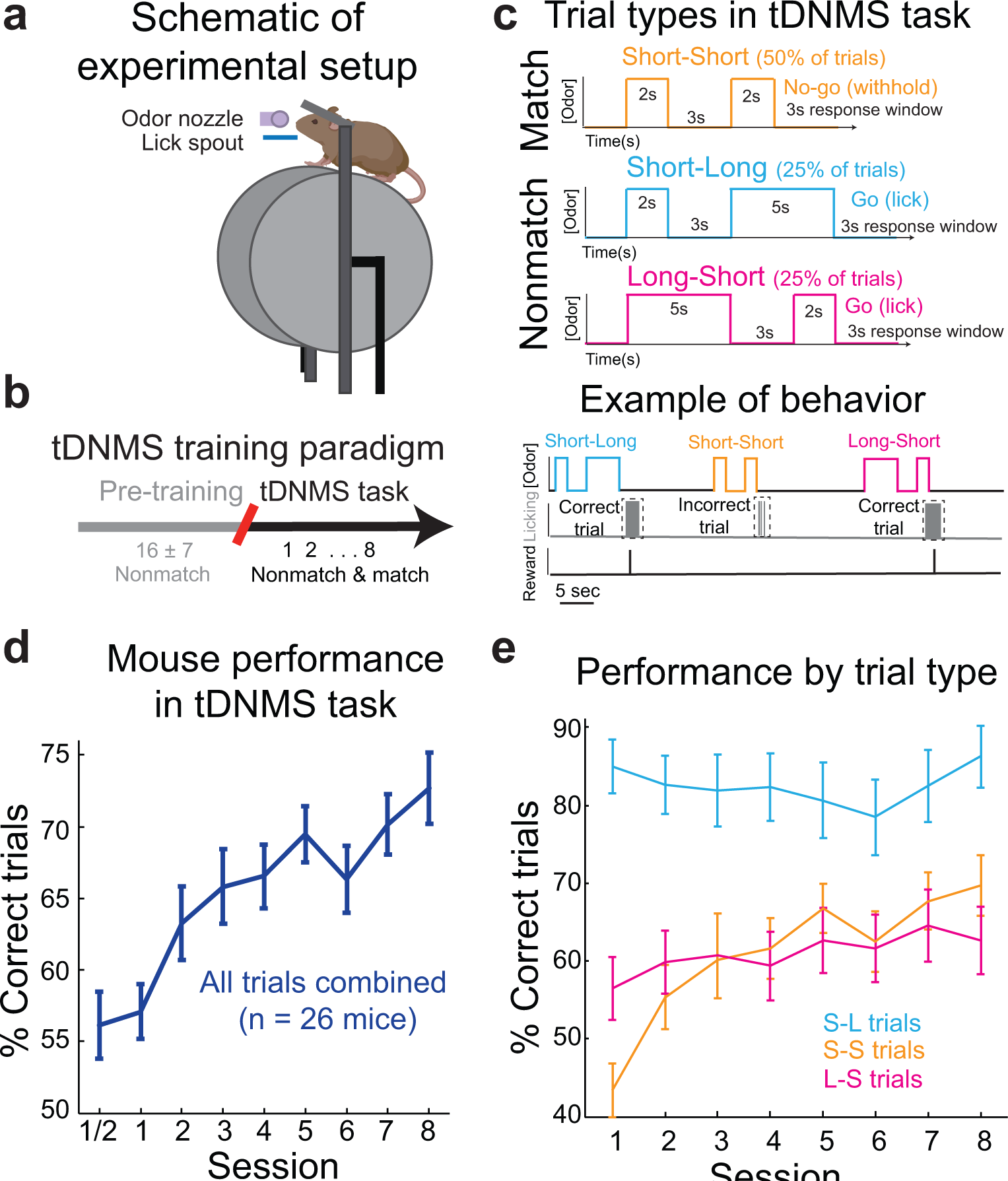
Mice learn a novel tDNMS task. **a.** Simplified experimental set-up. **b.** Overview of training paradigm. Mice are first pre-trained to lick at the offset of nonmatch trials to trigger reward delivery. Upon reaching criteria after an average of 16 sessions (mean ± s.d.), mice begin 8 training sessions on the tDNMS task, where match trials are introduced. **c.** Trial types and example behavior. The tDNMS task consists of 3 types of trials defined by stimulus durations: short-short (S-S), short-long (S-L), and long-short (L-S). To perform the task correctly, mice must lick in the response window following nonmatch trials and withhold licking in match trials. **d.** Percent correct across all trial types for each session, averaged across all mice (mean ± s.e.m.; n = 26 mice). **e.** Percent correct by trial type for each session, averaged across mice (mean ± s.e.m.; n = 26 mice).

Before beginning the tDNMS task, mice undergo a shaping procedure to learn the trial structure. Mice are only presented with nonmatch trials and learn to refrain from licking during the first odor and ISI of each trial, then lick near second odor offset to earn a reward (Fig. 1b, S1d; see Methods). After mice reach criteria (see Methods), indicating learning of the “odor, odor, response” trial structure, mice begin the tDNMS task, where match trials are first introduced and equally balanced with nonmatch trials over each session (90 trials/session over ∼45 mins). Notably, while mice could employ a simple strategy of licking after second odor offset during pretraining, the introduction of match trials transforms the task into one that requires timing. Therefore, in the tDNMS task, mice must learn to attend to the temporal structure of each trial to decide whether to lick or withhold.

To test whether mice learn the tDNMS task, we monitored behavioral performance in 26 mice over 8 training sessions on the tDNMS task (Fig. 1d). While mice began at chance performance on session 1, they steadily improved with training, averaging 73.16 ± 2.47% (mean ± s.e.m. for all data unless otherwise reported) correct responses on session 8, demonstrating learning (27.16 ± 4.13% change from session 1 to sessions 7&8; Repeated Measures ANOVA F_7,25_ = 8.43, p <0.001). To better understand the learning process, we examined performance by trial type. We expected mice to begin the task by licking at second odor offset, given their pretraining, and thus perform well in nonmatch trials. Indeed, mice averaged 85.35 ± 3.51 correct S-L trials on day 1 of the tDNMS task, compared to 91.05 ± 2.67% on the last half session of pretraining (Fig. 1e; Fig. S2a; Student’s Paired t-test, p = 0.24). Unexpectedly, performance on L-S trials dropped from 86.28 ± 2.01% correct responses on the last half session of shaping to 56.63 ± 4.1% correct on day 1 of the task (Fig. 1e; Fig. S2a; Student’s Paired t-test, p <0.001). Mice can miss nonmatch trials either by incorrectly withholding or by licking prematurely. Most mistakes were from premature licking, suggesting mice reverted to an impulsive action of licking after long odors previously observed in shaping (Fig. S2b-c). Match trials require learning a new response of withholding, so we expected learning to occur on S-S trials. Indeed, mice began with poor performance, averaging 43.39 ± 3.49% correct S-S trials on day 1 due to their tendency to lick near second odor offset (Fig. S2d), as if still applying an “odor, odor, lick” strategy learned in pretraining. However, by day 8, mice learned to withhold licking on match trials, reaching 69.95 ± 3.95% correct S-S trials (Repeated Measures ANOVA F_7,25_ = 4.81, p < 0.001). Together, our data demonstrate that mice learn the tDNMS task by learning to withhold licking selectively on S-S trials, as evidenced by 1) improvement on this trial type, 2) high performance on S-L trials, and 3) a tendency to miss L-S trials by licking early, not withholding licking.

Limiting the task to 3 trial types allowed mice to robustly learn the tDNMS task over 7-8 sessions, by which point performance begins to plateau (Fig. S12b: no significant improvement from extended training across sessions 8-14: Repeated Measures ANOVA F_6,10_= 0.45, p= 0.84). However, the task design with only three trial types could inadvertently lead mice to adopt a rigid behavioral strategy rather than a flexible cognitive strategy. For instance, mice might simply distinguish trials based on the total duration, thereby circumventing the need to assess the durations of individual stimuli. To address this, we conducted a series of control experiments to affirm that mice perform the task by attending to each odor duration. Initial experiments without odor established that olfactory cues are essential for task engagement, with a significant drop in performance on nonmatch trials without odor (2.25 ± 1.01% correct, see Fig. S1e). To determine if mice use a rigid strategy involving total trial duration, we trained a cohort of mice on a modified version of the tDNMS task and manipulated the ISI in a subset of probe trials to make nonmatch and match trials the same duration. The resulting performance across probe and standard trials was unaffected, suggesting that mice were indeed responding to individual stimuli and not the overall length of the trial (74.14 ± 6.71% correct on standard nonmatch trials versus 80.71 ± 6.02% correct on probe trials; Student’s Paired t-test, p = 0.19, see Fig. S1f). Importantly, our results do not confirm that mice compare durations; mice likely use a simpler strategy to solve the tDNMS task (discussed later). Nonetheless, any strategy they use requires them to 1) monitor stimuli durations and 2) make decisions based on the stimuli’s position within the trial structure. Therefore, the tDNMS task meets our goals of requiring mice to make flexible decisions based upon the temporal structure of each trial.

### MEC time cells fire in context-dependent trajectories during tDNMS task

To characterize the learning dynamics of MEC time cells, we applied methods we have previously developed for large-scale cellular resolution two-photon calcium imaging in MEC^20,26^ (Fig. 2a,b). We recorded from populations of layer II MEC neurons expressing GCaMP6s (see Methods; Fig. 2c-e, S3), across 6 mice (Field-of-view 430 ± 54 𝜇𝑚 medial to lateral by 380 ± 45 𝜇𝑚 dorsal to ventral; depth below the surface 105 ± 8 𝜇𝑚) as well-trained mice performed the tDNMS task (15 ± 4 days pretraining then 13 ± 8 days of tDNMS training to reach day N; 82 ± 5% correct trial performance). Across the total population (2056 active neurons), we found that 33.8% of cells exhibited regular time-locked activity at a particular moment in each trial (Fig. 2f, S4). Consistent with previous reports during interval timing behavior^20^, we found that different MEC “time cells” were selectively active at different delay times from the start of each trial, forming a regular temporal sequence that spanned the entire trial epoch (Fig. 2g, S5a).

**Figure 2.**
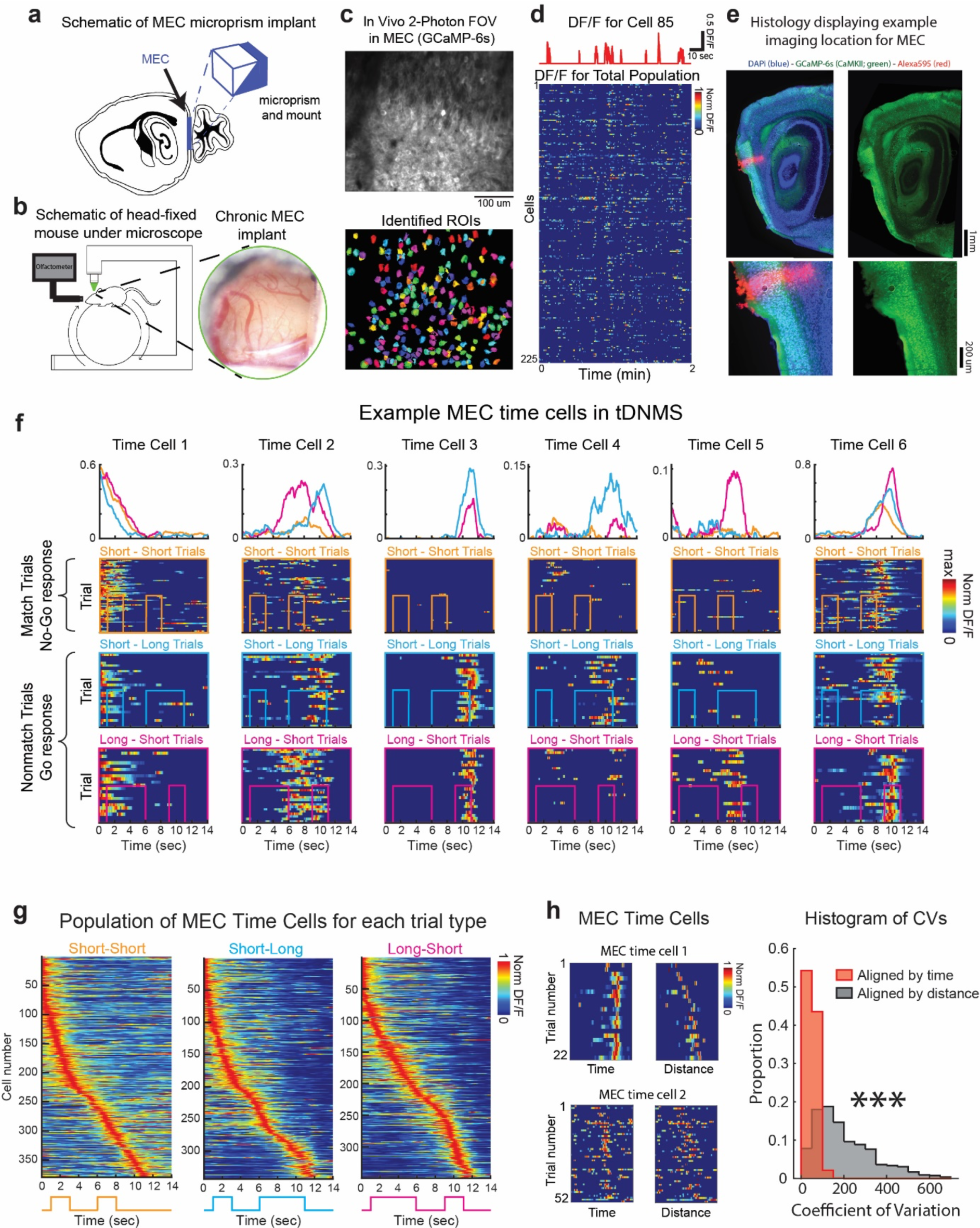
MEC time cells exhibit context-dependent sequential neural activity. **a.** Optical approach for imaging in MEC. **b.** Left, mouse positioned on the cylindrical treadmill and underneath the two-photon microscope. Olfactometer delivers odor near mouse snout. Right, image of MEC prep following chronic implant. **c.** Average image of example FOV labelled with GCaMP-6S (top) and ROIs from this FOV (below). **d.** Example dF/F time series for individual neuron (top) and all ROIs from panel c. **e.** Representative histology. Red pin mark indicates relative location of imaging field. **f.** Example individual MEC time cells. MEC time cell 1 is stable across all trial types (i.e. context-independent). MEC time cells 2-6 are significantly differentially active across specific trial types (i.e. context-dependent). **g.** All MEC time cells sorted for each trial type. **h.** Left, example time cells plotted as a function of elapsed time and distance travelled from trial onset. Right, distribution of coefficient of variation for all MEC time cells, measured as a function of elapsed time and distance travelled from trial onset. t(1070) = 39.9, p < 0.0001, n = 1,071, paired t-test.

During the tDNMS task, mice were free to run on a cylindrical treadmill. Since prior work has shown that many MEC neurons can encode distance travelled^27,28^, we wondered whether the time-locked activity of MEC time cells might be better explained by distance travelled from trial on-set. In support of the idea that MEC time cells encode elapsed time in the task, and not distance travelled, we found that the vast majority of time cells displayed a smaller coefficient of variation (CV) when measuring as a function of elapsed time versus distance travelled from trial onset (CV of elapsed time: 48.4 ± 0.8, elapsed distance: 205.7 ± 4.4, p < 0.0001, t_(1070)_ = 39.9,Paired t-test; Fig. 2h). Additionally, to estimate the specific contribution of different behavior variables (distance traveled (D), time elapsed (T) and licking (L)) on the activity of MEC neurons, we used a generalized linear model (GLM) to fit calcium activity (dF/F) as a Gaussian linear function of different combinations of the three behavior variables^29,30^. For the dF/F of each cell in each trial type, we fit 7 models which included 3 single variable models (D, T, L), 3 double variable models (TD, TL, DL), and 1 full model (TDL). We calculated the log-likelihood (LLH) gained by a specific variable as the difference between the full model and the reduced model. Consistent with results using the CV, our GLM results show that the log-likelihood gained by time is significantly greater than the log-likelihood gained by either distance or licking (Fig. S6), suggesting the vast majority of cells are tuned to elapsed time in the tDNMS task.

To perform the tDNMS task, subjects must perceive and use stimuli durations to determine trial type (i.e. nonmatch or match) and learn the appropriate response (i.e. Go or No-Go, respectively). Importantly, because trials consist of unique sequences of cues (differing only in duration) that dictate the appropriate behavioral response, we refer to each trial type as a temporal context. The robust learning and task structure allowed us to ask whether distinct populations of time cells encode each trial type, or temporal context. While some time cells showed stable activity across trial types (Time cell 1 in Fig. 2f), others displayed activity specific to the temporal context. This context-specific activity took various forms. Time field shifting (Time cell 2 in Fig. 2f) was rarely observed, and in many cases, time fields disappeared in response to changes in temporal context (Time cell 3∼5 in Fig. 2f). Additionally, some neurons modulated their firing rate according to the context (Time cell 6 in Fig. 2f). By examining time cell stability across each pair of trial types (S-S vs S-L, S-S vs L-S, S-L vs L-S; Fig. S5b), we found that more than half of the time cells exhibited stable time fields (58.4%, n = 944/1617). For those that “remapped” between contexts, time fields usually either disappeared in one trial type (33.2%, n = 537/1617) or changed activity level (5.32%, n = 86/1617). Few time cells shifted the timing of peak activity (3.09%, n = 50/1617). Since time field shifting was rare, the sequence of time cells sorted by time of peak activity remained coherent across trial types (Fig. S5a, c). Among all time cells, less than 20% of neurons (n = 132/695) remained stable across all context pairs, and the other 80% of neurons (n = 563/695) showed remapping at least one context pair. Therefore, our results demonstrate that unique populations of MEC time cells are used to represent distinct temporal contexts, forming a unique trajectory or “timeline” to represent each trial type.

### Context-dependent sequences support learning of flexible interval timing behavior

Initially, on day 1, mice do not utilize temporal context to guide behavior; however, over several training sessions, they learn to respond correctly on 70-90% of trials per session. If context-dependent MEC time cell activity supports task learning, our data should support several predictions. First, we would expect that context-dependent activity should be relatively absent on day 1 and should emerge over the course of learning. Second, if context-dependent MEC time cell activity supports learning of the correct response, then the coherence of individual time cell activity and/or the regular sequential neural activation across the population should be disrupted on “error trials”, when mice incorrectly report match or nonmatch.

To test the first prediction, we averaged the activity of MEC time cells for each trial type and compared the correlations for each cell’s response across trial types, before and after learning (see Methods). Because mice learn at varying rates, we classify sessions with over 70% correct trials as “day N” (training session 4-21). We found that the average correlation of time cell rate maps across match and nonmatch contexts is significantly lower on day N compared to day 1 (day 1: 0.57 ± 0.01, day N: 0.42 ± 0.02, p < 0.0001, z = 6.8, Wilcoxon rank sum test; Fig. 3a-c). These results demonstrate that context-dependent MEC time cell activity is more limited on day 1 and develops to become more distinct over the course of learning. During the tDNMS task, the information required to distinguish contexts accumulates throughout each trial, with key moments providing enough information for an “ideal observer” to discern the trial context. We wondered whether the population dynamics of context-dependent MEC time cells might be informative about this time-dependent decision process and provide further support that these context-dependent dynamics support learning in the tDNMS task. To test this, we measured the difference in dF/F for each time cell across trial types at successive moments in the trial epoch and averaged across time cells in order to generate population vectors. Compared to a null distribution with randomly assigned trial labels, we find that the neural dynamics significantly diverge from the null distribution at key moments after the stimuli have become distinct across trial context (Fig. 3d). Importantly, we find that over learning, these differences become more pronounced (Discriminant Index for S-S vs S-L: Day 1 = 0.40 ± 0.07, Day N = 0.67 ± 0.08, p = 0.01, z = 2.55; S-S vs L-S: Day 1 = 0.52 ± 0.07, Day N = 1.00 ± 0.09, p < 0.001, z = 3.80, Wilcoxon rank sum test; Fig. S5d). In the 3-trial structure, the long stimuli presented on L-S trials could be used to classify these nonmatch trials earlier in the trial epoch compared to the other trial types. Interestingly, we find that the neural dynamics diverge from the null distribution earlier when comparing L-S to S-S versus S-L to S-S trials (Fig. 3d). Using a separate measure to assess learning, we also find that from day 1 to day N, the population of MEC time cells shift on each trial type to encode later times in the trial, which correspond to moments when there is sufficient information to disambiguate the trial type (peak times on Day 1: 3.66 ± 0.17, Day N: 4.67 ± 0.18, p < 0.0001, z = 4.70, Wilcoxon rank sum test; early peak proportion on Day 1: 36 % (n = 164/450), Day N: 19 % (n = 76/402), p < 0.0001, χ^2^_(1)_ = 30.4, chi-squared test; Fig. 3e, f). Next, we asked whether the ensemble activity of time cells on individual trials contains enough information to accurately decode the temporal context. During the late phase of trials (9∼11 seconds), ensemble activity exhibited distinct separation according to trial type as shown by Linear Discriminant Analysis (LDA) (Fig. 3g, S7). To quantify this separation, we applied K-means clustering to each animal’s LDA plot, revealing a clustering accuracy substantially higher than the bootstrapped chance level in five out of six animals (For mouse 1 through 6: p= 0, 0.04, 0.08, 0, 0, 0.001, respectively). This result was further corroborated using an alternative classification method. Employing the Support Vector Machine (SVM), we constructed a model for classifying trials by type or as match vs. nonmatch (Fig. 3h). Interestingly, when the model was trained and tested using neural activity during the early phase of trials (0∼2 seconds), it failed to correctly identify trial information (Trial type decoding for mouse 1 through 6: all p-values > 0.1; Match vs Nonmatch decoding for mouse 1 through 6: all p-values > 0.1). In contrast, when utilizing activity during the late phase (9∼11 seconds), the model successfully decoded trial identity in most animals (Trial type decoding for mouse 1 through 6: p-value = 0, 0.002, 0.25, 0, 0, 0; Match vs Nonmatch decoding for mouse 1 through 6: p-value = 0, 0, 0.05, 0, 0, 0), aligning with the findings from the K-means clustering analysis. Together, these results support prediction 1 showing that context-dependent MEC time cell activity emerges over learning, eventually displaying an over-representation at later moments in the task, with large deviations in context-dependent neural trajectories near key moments in the task when it is possible to distinguish trial type.

**Figure 3.**
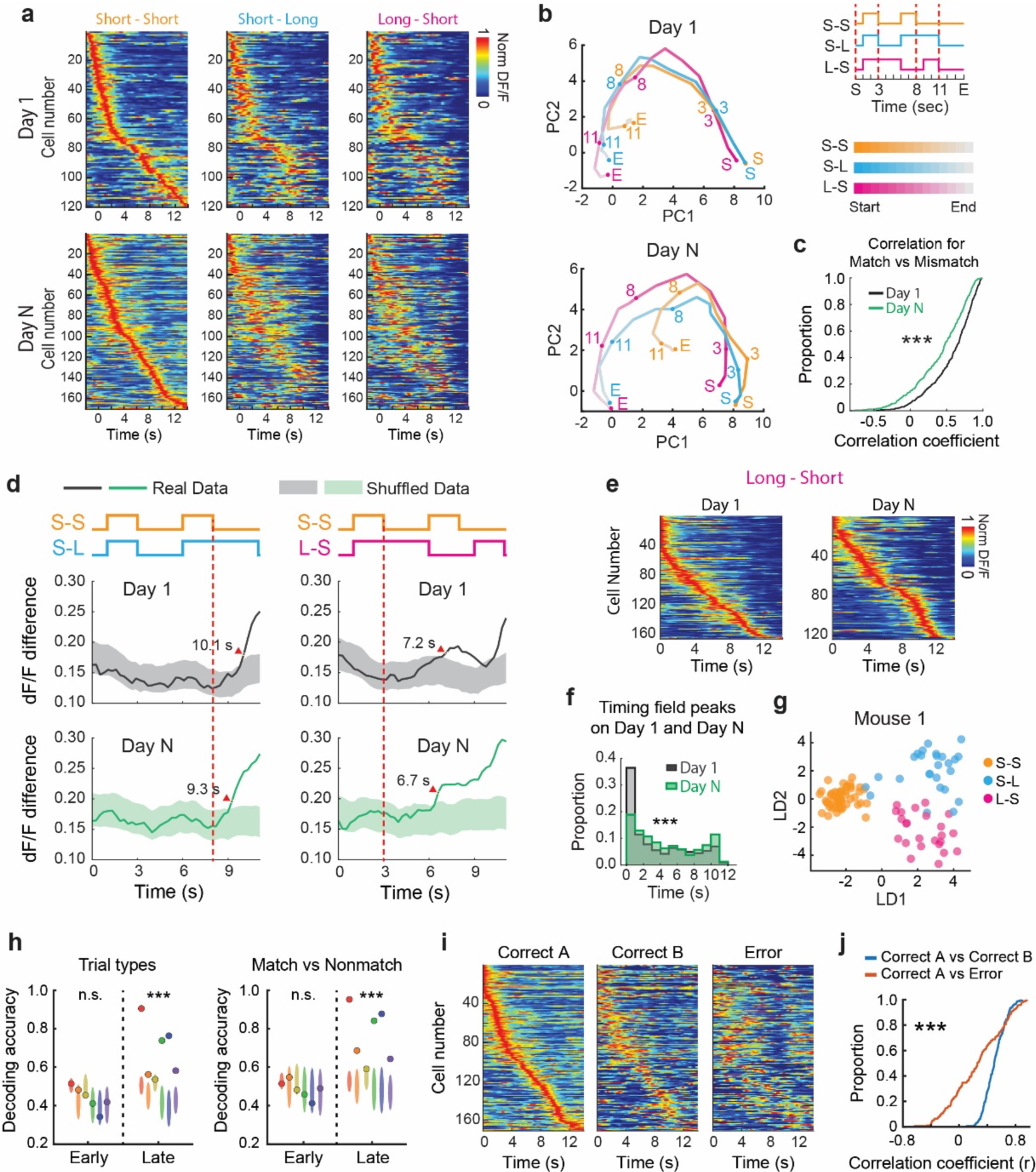
Context-dependent MEC time cell population dynamics support learning of flexible interval timing behavior. **a.** Top: The population of MEC time cells significantly tuned to Short-Short trials and sorted by response times during Short-Short trials, depicted for Short-Short (left), Short-Long (middle), and Long-Short (right) trials on day 1. Bottom: Same as top, but for day N sessions. **b.** Top two principal components displayed for population of MEC time cells across trial types. **c.** Mean Pearson’s Correlation Coefficients of MEC time cells across different trial types for day 1 (black) and day N (green), Day 1 n = 449, Day N n = 452, z = 6.8, p < 0.0001, Wilcoxon rank sum test. **d.** Left: Population vector for the difference in dF/F computed for each MEC time cell across SS and SL trial types as a function of trial time. Solid lines are the actual data, and shades indicate distributions of shuffle data. Right: The same for SS versus LS trials. **e.** Sorted sequence of MEC time cells in LS trials on day 1 (left) and day N (right). The box represents 25% percentile, median, and 75% percentile. **f.** Histogram of the timing of peak responses for all MEC time cells on day 1 (black) and day N (green), for all three trial types. Day 1 n = 450, Day N n = 402, z = 4.70, p < 0.0001, Wilcoxon rank sum test. **g.** LDA plots for individual trials for example mouse. **h.** Decoding accuracy of SVM models for trial types (left) or match vs nonmatch (right). The models are built on neural activity from either early phase or late phase of trials. Color circles depict the accuracy from actual data, with oval shades representing the chance distribution. Each color corresponds to an individual animal (n = 6). Trial type decoding in Early: z = 0.03, p = 0.97; in Late: z = 4.16, p < 0.0001; Match vs Nonmatch decoding in Early: z = 0.66, p = 0.51; in Late: z = 4.20, p < 0.0001, Wilcoxon rank sum test. **i.** MEC time cells during SS correct trials A (left; sorted by A), correct trials B (middle; sorted by A), and SS error trials (right; sorted by A) on day N of training in the tDNMS task. **j.** Cumulative distribution for Pearson’s correlation coefficients calculated for MEC time cells comparing correct trials A to B (blue), and correct trials A to error trials (red) (see Methods). Day 1 n = 120, Day N n = 170, z = 7.5, p < 0.0001, Wilcoxon signed-rank test.

To test our second prediction, we compared the average response for each MEC time cell for correct and error trials. The results show significantly higher coherence for time cells across randomly selected blocks of correct trials as compared to error trials on day N (correlation between Correct Trials - Group A and Correct Trials - Group B: 0.51 ± 0.01 Correct Trials - Group A and Error Trials: 0.26 ± 0.03, p < 0.0001, z = 7.5, Wilcoxon signed-rank test; Fig. 3i, j). Further, when comparing correct versus error trials on day 1 and day N, we find that the average correlation on day N is significantly reduced compared to day 1 (session type main effect: p < 0.0001, F_(1, 288)_ = 18.5; trial type main effect: p = 0.12, F_(1.93, 555.15)_ = 2.1; interaction: p = 0.33, F_(1.93, 555.15)_ = 1.1, two-way mixed ANOVA with trial type and session factors; Fig. S8), providing additional support that MEC time cell dynamics evolve over learning in the tDNMS task. Together, our results demonstrating 1) the emergence of context-dependent time cell activity over learning and 2) altered coding during error trials support the hypothesis that MEC time cells form unique trajectories used to encode the structure of each trial type and likely used to guide context-dependent timing behavior.

### Does MEC time cell activity flexibly adapt to changes in the temporal structure of the tDNMS task?

The observation of context-dependent neural dynamics suggests that the activity of MEC time cells can reflect the temporal structure of a trial. To expand upon this finding, we asked how time cells would adjust to manipulations in the trial structure. We recorded MEC time cells in the normal tDNMS task, then halfway through the recording session introduced a lengthened ISI of 5 seconds (Fig. S9a). Mice responded to the change in the ISI by delaying approach behavior and predictive licking (Fig. S9b). We quantified this through the center of mass of velocity and licking (in units of seconds) on normal and probe nonmatch trials: velocity in S-L trial type: normal (4.73 ± 0.11 s) vs probe (5.67 ± 0.10 s), p < 0.0001, z = 5.41; velocity in L-S trial type: normal (4.92 ± 0.19 s) vs probe (5.20 ± 0.10 s), p = 0.0065, z = 2.72; licking in S-L trial type: normal (13.08 ± 0.07 s) vs probe (14.51 ± 0.06 s), p < 0.0001, z = 8.72; licking in L-S trial type: normal (13.12 ± 0.08 s) vs probe (14.47 ± 0.03 s), p < 0.0001, z = 8.07; Wilcoxon rank sum test), indicating mice perceived the difference in the trial structure during the probe trials. Notably, on S-S trials mice did not exhibit clear approach behavior or licking, making it difficult to measure the effects of S-S probe trials. We found that, on average, population-level MEC time cell activity was delayed in response to the longer ISI (peak times on normal and probe trials: S-S trial types: normal (6.3 ± 0.4 s) vs probe (7.8 ± 0.5 s), p < 0.0001 z = 3.9; S-L trial types: normal (4.3 ± 0.5 s) vs probe (6.2 ± 0.6 s), p < 0.0001, z = 4.5; L-S trial types: normal (5.1 ± 0.6 s) vs probe (7.5 ± 0.6 s), p < 0.0001, z = 4.2; Wilcoxon signed-rank; Fig. S9c, d). These results suggest that populations of MEC time cells can flexibly adapt to trial structure, incorporating the animal’s prior of the relative duration of the stimuli and ISI.

### MEC is required to learn flexible, context-dependent timing behavior

The emergence of context-dependent MEC time cells provides a potential neural dynamical mechanism that could underlie tDNMS learning, where the formation of context-specific “timelines” (i.e. time cell trajectories) could allow animals to differentiate trial types. To causally test if MEC activity is necessary to learn the tDNMS task, we used a chemogenetic approach to inhibit MEC during task learning (Fig. 4a). We first validated our approach by local injection of AAV expressing the pharmacologically selective designer G_i_-coupled muscarinic receptor (hM4D) along the dorsal-ventral extent of MEC in transgenic mice with GCaMP6s expression under the CaMKIIa promoter (Fig. S10a; see Methods). Using in vivo two-photon Ca^2+^ imaging, we observed that the frequency of Ca^2+^ transients was reduced by 80% at 30 minutes post 1mg/kg DCZ injection compared to baseline (n=302 neurons across 2 mice; KS test, p < 0.01), confirming our ability to inhibit MEC (Fig. S10b).

**Figure 4.**
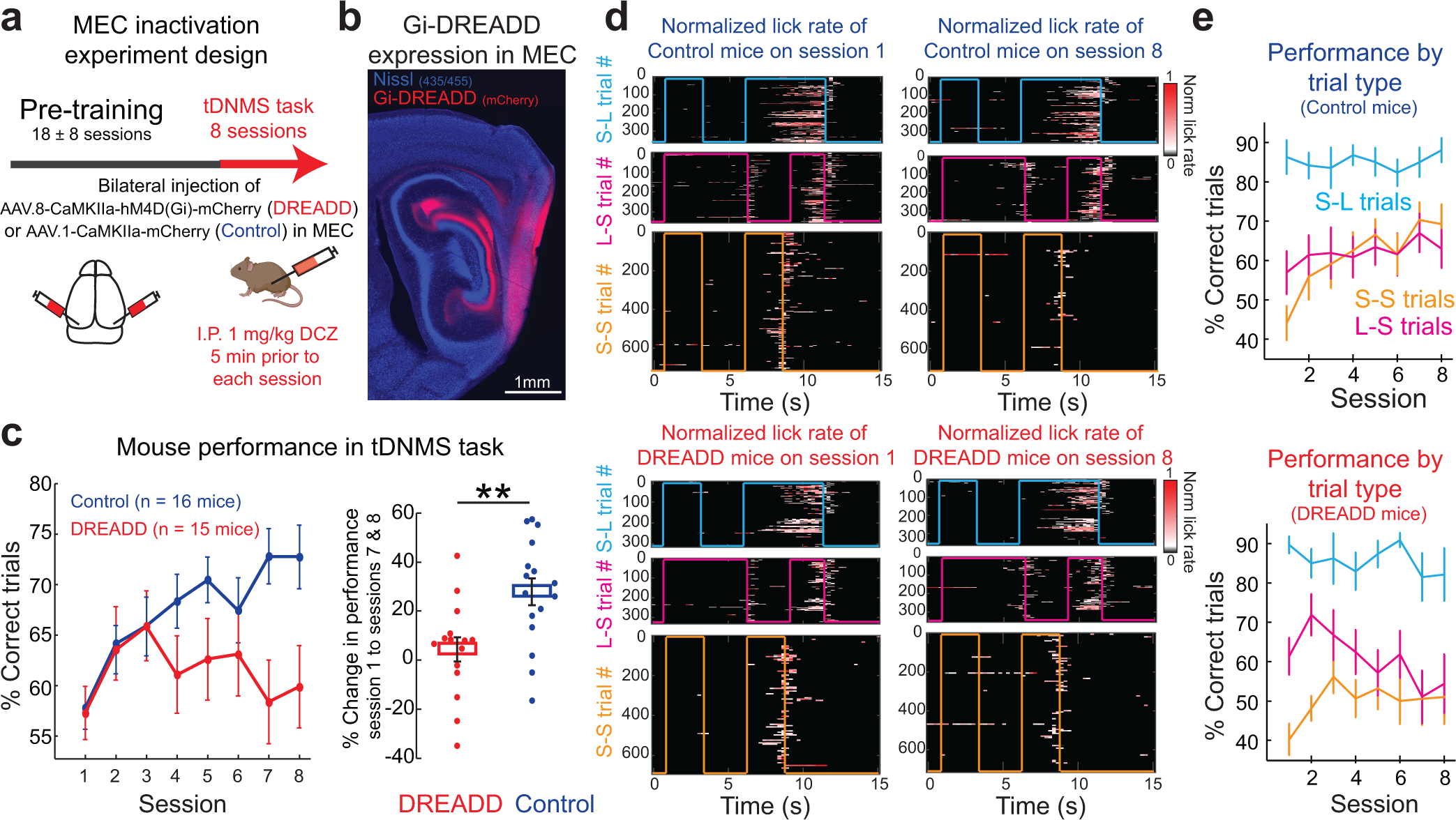
MEC is necessary to learn the tDNMS task. **a.** Experimental paradigm. Mice received bilateral injections of either an inhibitory DREADD (n=15) or control virus (n=16). Mice began pre-training with only nonmatch trials and learned to lick at the offset of second odor stimuli. Following pre-training, mice began the tDNMS task. DREADD agonist DCZ (1mg/kg) was administered via I.P. 5 minutes prior to each of the 8 tDNMS sessions. Pretraining sessions reported as mean ± s.d. **b.** Sagittal section depicting hM4D(Gi)-mCherry expression in MEC with blue Nissl staining. **c.** Left, performance on the tDNMS task for Control (blue) and DREADD (red) mice. Right, percent change in correct responses from day 1 to average of days 7 and 8 (p<0.01, unpaired t-test). **d.** Top, licking behavior during tDNMS task for Control mice on session 1 (left) and session 8 (right) for all 3 trial types. Bottom, licking behavior during tDNMS task for DREADD mice on session 1 (left) and session 8 (right) for all 3 trial types. Consummatory licking following reward delivery is not shown. **e.** Average performance for Control (top) and DREADD mice (bottom) on each of the three trial types. Bars indicate mean performance ± s.e.m calculated across mice.

To test whether MEC activity is necessary for tDNMS learning, we bilaterally injected AAV expressing hM4D across the dorsal-ventral extent of MEC (Fig. 4b; Fig. S13). Mice then underwent water restriction and shaping before beginning the tDNMS task. We monitored learning across 8 sessions of the tDNMS task and administered the DREADD agonist DCZ via I.P. injection 5 minutes prior to each session to inhibit MEC for the duration of training. As expected, control mice learned the task within 8 days, averaging 72.76 ± 2.62% correct responses across sessions 7-8 (Fig. 4c) (Repeated Measures ANOVA F_7,15_ = 5.09, p < 0.001). In contrast, DREADD mice showed no improvement from session 1 (54.07 ± 3.79% correct responses) to sessions 7-8 (59.10 ± 3.28% correct responses) (Repeated Measures ANOVA F_7,14_= 1.18, p= 0.32). Inactivation of MEC prevented mice from learning the tDNMS task.

To further investigate the specific deficit caused by MEC inhibition, we analyzed data by trial type. We found that both control and DREADD mice performed well on S-L trials, starting immediately on session 1 (Fig. 4e). Further analysis indicated this is likely supported by learning that took place during the shaping phase of the task (before DCZ injections) (Fig. S11a). On Day 1, both control and DREADD mice exhibited poor performance on L-S nonmatch trials. This was likely a result of impulsive tendencies causing the mice to lick at the end of the long odor presentation, a behavior noted during the shaping phase (see Fig. S11b-c). Since match trials uniquely demand the learning of a new withholding response, we anticipated that improvement in these trials would drive overall learning. In line with this, control mice demonstrated a significant increase in S-S trial accuracy across 8 sessions (from 44.16 ± 4.34% correct on session 1 to 69.27 ± 5.01% correct on session 8; Repeated Measures ANOVA F_7,15_ = 5.93, p < 0.001, as shown in Fig. 4e). Interestingly, DREADD mice showed an initial improvement in S-S trials over sessions 1-3 (40.43 ± 4 % correct match trials on session 1 vs. 56.31 ± 4.17% correct match trials on session 3; Repeated Measures ANOVA F_2,14_ = 8.01, p <0.01), reflected in their overall learning curve. However, match trial performance did not significantly improve across the 8 sessions (Repeated Measures ANOVA F_7,14_ = 0.78, p = 0. 61), leveling near 50% (50.69 ± 3.63% correct match trials over sessions 6-8), indicating that mice guessed whether to lick or not. While control mice learned to use stimulus durations to determine when to withhold licking, DREADD mice perseverated with the rigid strategy of licking at trial offset (KS test on Control and DREADD S-S learning curves, p < 0.01) (Fig. 4d). This pattern suggests that MEC inhibition impaired the ability to form memories based on the new temporal context introduced on day 1 in the tDNMS task.

Notably, DREADD mice were not impaired in all aspects of timing behavior. Both control and DREADD mice performed above chance in nonmatch trials suggesting that MEC is not required to recall well-learned temporal contexts like “S-L-reward” (Fig. 4e). Mice even engaged in predictive licking in anticipation of reward (Fig. 4d), further indicating that MEC is not required to perceive or estimate learned durations. The key behavioral difference between control and DREADD mice was that DREADD mice were unable to form a memory of a new temporal structure, leading to an inability to adopt a flexible context-based strategy to solve the task.

An alternative explanation for our results is that MEC inhibition may not affect the learning of temporal context, but rather, other aspects of behavior necessary for tDNMS performance. We therefore performed additional analyses to determine whether MEC inactivation impacted non-timing behavior. Initially, we considered whether MEC inactivation could impair odor perception. However, the robust performance by DREADD mice on nonmatch trials clearly indicate intact odor perception (Fig. 4d-e). We also examined whether MEC inhibition might increase impulsivity, as success in the task requires mice to inhibit licking. Should impulsivity be heightened, mice would find it challenging to withhold licking even with an understanding of the task. To investigate potential differences in impulsivity between control and DREADD mice, we compared the average time from trial onset to first lick for both control and DREADD mice across session 1. If MEC inhibition caused mice to be more impulsive, DREADD mice should lick earlier. This was not the case, indicating MEC inhibition did not make mice more impulsive (Fig. S10c). Thus, our data confirm that MEC inhibition causes a specific deficit in learning context-based timing behavior.

### MEC is not necessary for ongoing tDNMS performance

The emergence of context-dependent time cells led us to test, and confirm, the hypothesis that MEC is necessary to learn the tDNMS task. We next wondered whether the role of MEC is confined to learning, or whether MEC is also required for ongoing task performance. To distinguish these possibilities, we performed an additional experiment in which we silenced MEC after task learning. We did this as a continuation of our first MEC inactivation experiment: following 8 consecutive sessions of MEC inactivation (shown in Fig. 4), we took mice off DCZ and instead administered saline over sessions 9-14 (Fig. S12a), hypothesizing that without MEC inhibition, mice would learn the task. Indeed, without MEC inhibition, DREADD mice learned the task (Repeated Measures ANOVA F_5,8_ = 3.72, p <0.01), reaching 73.65 ± 2.9% correct responses over sessions 13-14 (Fig. S12b). Because mice successfully learned, we returned to DCZ to inhibit MEC over sessions 15-16 (Fig. S12a-b). Mice performed well on sessions 15-16 despite MEC inhibition: there was no difference in behavior from sessions 13-14 to 15-16 (2.19 ± 7.30% change in correct response from sessions 13&14 to sessions 15&16 for DREADD mice, n = 7), showing that MEC is not required to perform the tDNMS task after learning (Fig. S12b). These results suggest that MEC is specifically necessary to form representations of temporal context; other brain regions can guide post-learning performance.

### MEC is not required to learn rigid timing behavior

A key requirement in developing our timing task was cognitive flexibility, given findings that MTL structures are involved in flexible, not rigid navigation behavior^4,6,31^. The critical role of MEC in learning the tDNMS task therefore led us to question whether MEC is necessary to learn any temporal relationship, or whether MEC is specifically required for flexible tasks requiring distinguishing between temporal contexts. To disentangle these possibilities, we trained DREADD and control mice on a simple, rigid, fixed interval (FI) task, in which a drop of water was delivered to the head-fixed mouse every 10s^32^. Predictive licking, which increases prior to the reward, signals an understanding of the task’s temporal structure (Fig. S12c). To test if MEC is needed to learn this simple timing task, we administered DCZ prior to each of 5 training sessions to inactivate MEC. To assess learning, we calculated the percent of trials in which mice engaged in predictive licking. Over 5 sessions, the percent of trials with predictive licking increased for both control (16.12 ± 5.34% on day 1 vs 57.76 ± 7.27% on day 5; Repeated Measures ANOVA F_4,10_ = 7.05, p < 0.001) and DREADD (14.67 ± 2.42 % on day 1 vs 70.88 ± 4.39 on day 5; Repeated Measures ANOVA F_4,9_ = 36.84; p <0.001) mice, indicating mice learned to anticipate the forthcoming reward (Fig. S12d-e). Additionally, as the sessions progressed, the peak of licking activity for both groups shifted closer to the reward time, suggesting an improved precision in timing (control: -2.05 ± 0.12s on day 1 to -0.96 ± 0.16s on day 5; Repeated Measures ANOVA F_4,10_ = 9.20; p < 0.001; DREADD: -1.73 ± 0.14s on day 1 to -0.97 ± 0.11s on day 5; Repeated Measures ANOVA F_4,9_= 7.75, p < 0.001) (Fig. S12f). There was no significant effect of experimental condition on learning as measured though either the percent of trials with predictive licking (Repeated Measures ANOVA group x time F= 1.98, p = 0.11; KS test on learning curves, p = 0.7) or time of peak licking (Repeated Measures ANOVA group x time F = 1.05 p = 0.39; Unpaired t-test on day 5 peak lick times, p = 0.99), indicating that MEC inhibition did not affect learning. Therefore, MEC is not needed to learn rigid temporal relationships, indicating a specificity in learning context-dependent timing behavior.

### The use of cognitive strategies to solve the tDNMS task

As previously mentioned, the tDNMS task can be solved through multiple strategies, each requiring a varying degree of cognitive flexibility. Therefore, we sought to 1) identify the strategies utilized by mice and 2) determine the involvement of MEC in relation to the degree of cognitive flexibility, thereby providing additional evidence of MEC’s role in learning flexible timing behavior.

We delineated four “cognitive strategies” which mice could use to solve the tDNMS task: Strategy 1 – animals identify the long duration odor as a “go” cue then wait for the second odor offset to respond (does not necessarily require timing both odors); Strategy 2 – animals time the entire trial duration (through the end of the second odor) and respond on longer trials; Strategy 3 – similar to 1, mice use the long stimulus as a go cue, but still time the duration of both stimuli (permitting predictive licking prior to the second odor offset); Strategy 4 – mice observe both odor cues then perform an explicit comparison of the durations. Importantly, these 4 cognitive strategies require recognition of the odor-odor-response trial structure. In contrast to these strategy-based approaches, mice might employ a simple “cue-based” approach, by learning to lick in response to specific events within a trial without fully grasping its structure—altering the likelihood of licking following any odor offset or specifically after the second odor or any prolonged odor cue, without integrating this into a two-odor-response framework. While this probabilistic approach may not always be successful, it can still result in high performance when fine-tuned.

Hallmarks for the use of each strategy would be present in behavior, both in the anticipatory licking and errors made. Therefore, to evaluate which strategy a given mouse is using, we modeled the observed lick counts as a non-homogeneous Poisson process based on weighted combinations of both cue-based features and each of the cognitive strategy-based features (Fig. 5a,b, S14a,b,e). We find evidence that individual mice may be using different approaches to the task, but most mice exhibit behavior that fits better to models that include a strategy-based feature (Fig. 5c, S14c). Accordingly, incorporating strategy-based features led to an improvement in model fit on correct as opposed to incorrect trials across all trial types, as well as on trials where mice had to actively lick (L-S or S-L) as opposed to withhold licking (S-S, Fig. S14d). Further, we found that including Strategy 2 or 3 resulted in the largest increase in model fits to the data (Fig. 5d). To distinguish between these two models, we investigated probe trials with manipulated ISIs (Fig. S1f, S9). By training our model on the standard ISI trials and testing on the subset with either shortened or lengthened ISI, we find that animals are unlikely to employ Strategy 2 (Fig. 5e). Interestingly, some mice perform the task fairly well employing an approach that incorporates only lick responses to the cue-based features: while model fit is significantly correlated with task performance (linear regression r=0.369, p=0.045), the improvement by adding in strategy-based features is not (r=0.025, p=0.895, Fig. S14f,g). This finding indicates that some mice might solve the task through altering lick probability in response to odor cues without necessarily using an odor-odor-response strategy.

**Figure 5.**
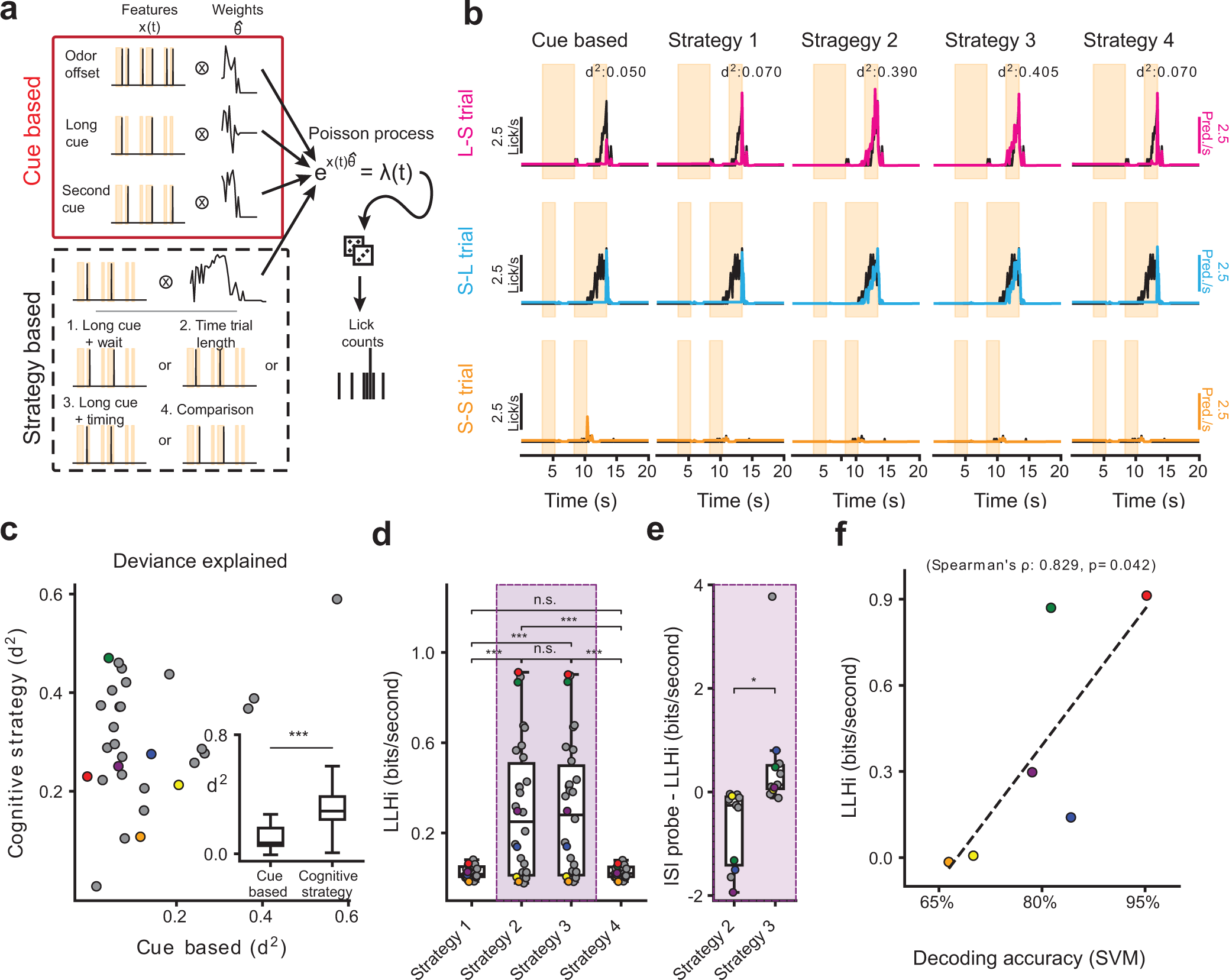
Model of behavior demonstrates potential cognitive strategy at use in tDNMS task. **a**. Model of the animals’ behavior as a non-homogeneous Poisson process. Expected lick counts are based on both event-based cues (odor offset, long odor offset, and second cue offset), as well as a cognitive strategy which could improve task performance but requires timing longer or multiple durations. For *x(t),* lags not shown. **b**. Comparison of cue-based and several potential strategy-based models for one mouse. Mean lick rate for each trial type shown in black, model predictions overlaid in color. **c**. Adding strategy-based features improves model fit to behavior, on average, over only cue-based features. Points from highest performing strategy-based model shown. *Inset.* Mean deviance explained (d^2^) is higher for strategy-based models (n=30 mice, p<0.001, paired t-test). **d**. Comparison of the log-likelihood increase (LLHi) over a cue-based model for the 4 different potential strategies (One-way ANOVA: F=16.57, p<0.001; Tukey test post-hoc test for multiple comparison of means: * - p<0.05, ** - p<0.01, *** - p<0.001, n.s. - not significant.) **e**. Testing the model on probe-trials with manipulated ISI times shows that Strategy 3 is more likely than Strategy 2, (p=0.03, paired t-test). **f**. Tendency to use more abstract methods during the tDNMS task is correlated with neural dynamics in MEC, based on best fit strategy and trial-type decoding results (ρ=0.83, p=0.04, Spearman’s rank correlation coefficient). Colors in panels **c**-**f** indicate imaged mice, gray indicates not imaged.

Our modeling also allowed for probing the neural dynamics in MEC time cells to predict the strategies used by each animal. Interestingly, trial-type decoding performance correlates with the use of a strategy-based approach (Spearman’s rank correlation coefficient, ρ=0.829, p=0.042, Fig. 5f). In contrast, no significant relationship exists between decoding accuracy and model fit in general when also considering the cue-based model (ρ=0.600, p=0.208, Fig. S14h). Together, our modeling results suggest that individual mice may apply different methods to solve the tDNMS task, with most utilizing Strategy 3, which involves using the long stimulus as a go cue while timing both stimuli. Furthermore, these results indicate that strategies requiring an understanding of the task structure and temporal context likely engage MEC more intensively.

### MEC time cells display coherent phasic activity during task and non-task epochs

This study was inspired from drawing parallels between mechanisms of spatial and temporal coding within MTL structures. There has been a growing body of evidence that a continuous attractor network (CAN), mediated by local recurrent synaptic connectivity in MEC, drives the neural dynamics of spatially selective grid cells^33–36^. Given the strong experimental support for this model in MEC, we wondered whether MEC time cells might also be driven through the same CAN mechanism. In this model, structured recurrent synaptic connectivity drives an “activity bump” in a local subpopulation of neurons. This activity bump is then translated across the network as a function of feedforward input which results in regular phasic activity among neurons in the CAN. A strong prediction of this model is that the relative phasic activity of cells in the network should be coherent during task and non-task relevant epochs. To test this prediction of the CAN model, we first measured the pairwise correlations of MEC time cell activity during the tDNMS trial epoch and compared these to the pairwise correlations when mice were not actively timing during the ITI (Fig. 6a). Both on day 1 and day N of recording, we find that the coherence of pairwise activity between trial period and inter-trial interval (ITI) is strongly positively correlated and much higher than the chance level (Day 1: Pearson’s r = 0.81 in actual, r = 0.29 in shuffle, p < 0.0001, z = 62.65; Day N: Pearson’s r = 0.76 in actual, r = 0.27 in shuffle, p < 0.0001, z = 64.57; Fig. 6b). Next, we sorted MEC time cells according to their relative phases during the tDNMS task and computed the pairwise correlation between MEC time cells during the ITI as a function of the time difference in the peaks of their firing fields in the tDNMS task. Our findings reveal that pairs of MEC time cells active at similar times during the tDNMS task also exhibit a high likelihood of concurrent firing during non-task periods (Pearson’s r = -0.12, p < 0.0001; Fig. 6c). We further repeated the same analysis for correct vs error trials (Fig. S15a) and for different trial types (Fig. S15b). In all comparisons, we found that time cells exhibited coherent activity across conditions (all p-values < 0.0001). These results are consistent with key predictions of a local recurrent CAN model that may support the regular, sequential activity of MEC time cells during timing behavior, suggesting both MEC time and spatial coding neurons could be driven through a common local circuit mechanism.

**Figure 6.**
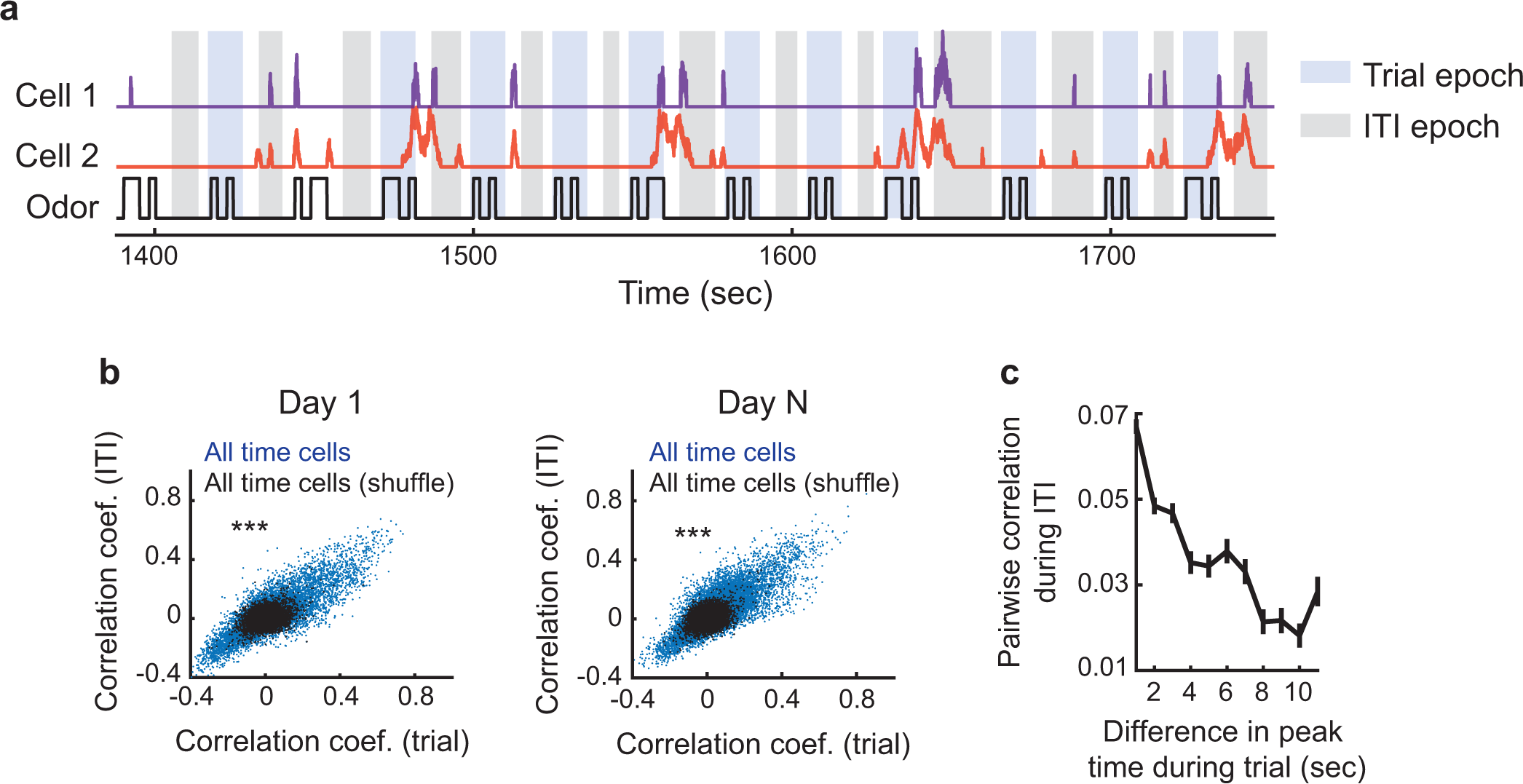
Activity correlation between time cells maintained across task and non-task epochs. **a.** example time series display for pair of cells during task and during the ITI. Odor stimuli shown below. **b.** Left, pairwise correlation between all time cells during the tDNMS task and during the ITI for real (blue) and shuffled data (black) for day 1 recording sessions. Right, same as in b, except shown for day N recording sessions. Day 1: n = 11,594, z = 64.57, p < 0.0001; Day N: n = 15,727, z = 64.57, p < 0.0001, Chi-squared test. **c.** Pairwise correlation of MEC time cells during the ITI as a function of their relative peak timing field activity during the tDNMS task. (mean ± s.e.m.)

## Discussion

A system capable of episodic memory must be able to track the duration of and between events to form an accurate memory of how experiences unfold over time. However, it remains unclear how MTL structures involved in episodic memory represent temporal relationships of events occurring on the order of interval time (seconds-minutes). Drawing inspiration from spatial literature, we hypothesized that distinct populations of time-selective neurons could form distinct “timelines” to encode the temporal structure of different experiences (temporal context). We tested whether previously identified MEC time cells form such “timelines” by examining their learning dynamics, testing the predictions that 1) distinct patterns of time cells should emerge as animals learn to distinguish temporal contexts, and 2) formation of such trajectories should be critical for learning temporal relationships.

Addressing our hypothesis required a multifaceted approach. We first developed a novel tDNMS paradigm that requires mice to differentiate the temporal structure of trials. By applying the tDNMS task in tandem with in vivo neurophysiological recordings, we confirmed our ability to record MEC time cells. We demonstrate that cells are tuned to elapsed time and not other features of behavior. We then leveraged the tDNMS task design to ask whether different trial types, or temporal contexts, are represented by different populations of time cells, as hypothesized. Our results show that distinct populations of time cells are active on distinct trial types, demonstrating that MEC time cells from unique “timelines” of each context. Crucially, multiple lines of evidence link the observed “timelines” to a role in learning of timing behavior. First, we find that populations of time cells exhibit distinct context-dependent neural dynamic trajectories, which diverge from a common trajectory, at key moments in the task when there is sufficient information to disambiguate the trial structure. Second, we find these trajectories become more distinct over learning across sessions. Third, we show that over learning, the population of MEC time cells shifts to over-represent later times in the trial, when the animal can disambiguate trial type and thereby solve the instrumental task. Fourth, on error trials, when the animal mistakenly classifies the trial context, we find that the regular sequence of MEC time cells is disrupted. Finally, we find that MEC inactivation prevents tDNMS learning, specifically by preventing the learning of a new temporal context. Combined, our results indicate that MEC time cells form distinct representations of temporal context, enabling animals to flexibly learn the temporal structure of experiences.

Interestingly, we find that MEC is not always required for learning and memory of interval timing behavior. MEC is not necessary to learn simple durations, reproduce previously learned durations, or recall learned contexts. Rather, MEC is selectively required to learn flexible, context-dependent temporal relationships. This finding fits our physiology data, implying that the role of MEC is to form unique representations of temporal context needed to support such flexible behavior. In further support of this idea, our computational modeling work demonstrates that mice employing a cognitive strategy show increased decoding accuracy of the trial context from the activity of MEC time cells. This specificity of MEC in flexible timing behavior is reminiscent of the finding that MTL structures play a key role in flexible, but not rigid forms of spatial navigation behavior^21^. Just as multiple memory systems guide navigation^31^, we suspect multiple memory systems guide timing behavior. Prior work has implicated basal ganglia and striatum, frontal, and parietal regions in timing^32,37–43^, providing evidence of other neural “clocks” that likely drive other aspects of timed behavior. Determining the constraints under which distinct clocks drive behavior, and interactions between clocks, will be an important step in understanding how the brain performs interval timing.

Prior studies investigating the role of MEC in interval timing have come to somewhat different conclusions about the contribution of MEC. Our observation that MEC is not needed to learn or time fixed intervals appears to contrast prior work describing roles of MEC in a) learning to remain immobile for a fixed duration^17^, b) precisely timing a learned duration^19^, and c) delay-dependent timing^18^. To reconcile these differences, we argue that each task involves an element of flexibility, such as a) updating behavior from a different duration used during pretraining, i.e. learning a new temporal context, b) updating a reference memory after failed trials, i.e. updating a memory of temporal context, and c) making delay-dependent associations, which requires learning temporal context. We expect that the flexibility of learning or updating a memory of temporal context requires MEC, bridging the results of each study. While our study focused on MEC, other MTL structures including the hippocampus and lateral entorhinal cortex also encode time. Prior work has examined temporal coding in various ways, including through explicit timing behavior^44,45^, sequence coding^46,47^, the delay period of tasks^48,49^, and timescales spanning minutes to hours to days^50–52^ . Many of these processes occur in parallel, making it difficult to pinpoint precise neural dynamics involved in each aspect of temporal coding. Our study focused on one aspect of temporal coding-interval timing. Given the clear role of MEC in our tDNMS task, a clear future direction will involve testing the necessity of other MTL regions in this task.

Our rationale for examining the role of MEC in interval timing, rather than other MTL structures, stemmed from drawing parallels between spatial and temporal coding. Since the discovery of spatially selective MEC grid cells there has been a strong focus on the role of MEC in navigation and spatial memory, leading to a research program that has given substantial insight into the circuit mechanisms that underlie grid cell firing. Namely, there is strong support for a continuous attractor network (CAN) that is mediated through structured local recurrent synaptic connectivity of MEC neurons^33–36^. A key feature of this model is that the network integrates synaptic input coding for animal heading direction and velocity, thereby driving sequential activity of a population of grid cells, each coding for different spatial phase(s) within an environment. This process is mathematically equivalent to path integration^33,53^, giving rise to a measure of distance travelled from a start location. We find this computation conspicuously similar to that of a clock, which rather than a measure of distance, can integrate a constant input to give rise to a measure of duration from a start time. A strong prediction of the CAN model is that neurons encoding similar phases while engaged in a relevant behavioral task, such as firing at similar locations in space during navigation or similar delay times during interval timing, will display coherent phasic activity during non-task relevant epochs such as the ITI. Consistent with this prediction, we find that the correlational structure of MEC time cells, as defined during the tDNMS task, remains coherent during the intertrial interval when there is no timing demand. We suspect that during timing, the sequential activity of MEC time cells is driven through similar continuous attractor network dynamics, that may have evolved for similar and often overlapping navigation processes across time and space. Accordingly, our findings suggest MEC neurons may serve as a general integration circuit, calculating either distance or time based upon relative behavioral demands.

## Materials and Methods

### Surgery and Behavior

#### AAV injections

All experiments were approved and conducted in accordance with the University of Utah Animal Care and Use Committee. To inhibit MEC, C57-BL6 mice (n = 40 mice: 20 male & 20 female; postnatal 2-3 months) were injected bilaterally with pAAV-CaMKIIa-hM4D(Gi)-mCherry (Addgene: AAV8; 2.40 x 10^13^ GC/mL; diluted 1:1 in PBS) or pAAV-CaMKIIa-mCherry (Addgene: AAV1; 1.40E x 10^13^ GC/mL; diluted 1:1 in PBS). A Nanoject III Injector (Drummond) was used to inject 80nL of virus (divided into 4×20nL injections, injected at a rate of 10 nL/s) at 6 sites in each hemisphere. Injections were targeted at 2.9mm lateral from bregma and 0.15mm rostral of the transverse sinus; at 3.3mm lateral from bregma and 0.15mm rostral of the transverse sinus; and at 3 depths (1.2mm, 1.6mm, and 2mm) from the dorsal surface of the brain.

#### MEC microprism implant

Methods for MEC microprism implant have been described previously^20,26^. Briefly, 6 (5m and 1f) C57-BL6 mice (∼P70) were anesthetized using 1–2% isoflurane. An approximately rectangular craniotomy was made over the dorsal surface of the cortex (above MEC) and cerebellum with corners positioned as follows: (i) ∼2.1 mm lateral of bregma, ∼4.5 mm caudal of bregma (∼300–500 µm rostral of the transverse sinus); (ii) ∼4.5 mm lateral of bregma, ∼4.5 mm caudal of bregma (∼300–500 µm rostral of the transvers sinus); (iii) ∼2.1 mm lateral of bregma, ∼7.75–8 mm caudal of bregma (∼3.25–3.5 mm caudal of the transverse sinus); and (iv) ∼4.5 mm lateral of bregma, ∼7.75–8 mm caudal of bregma (∼3.25– 3.5 mm caudal of the transverse sinus). After the skull was removed, a portion of the cerebellum was aspirated to expose the caudal surface of the cortex. The tentorium separating the cerebellum and cortex was carefully removed, leaving the dura of the cortex completely intact. A microprism (right-angle prism with 1.5-mm side length and reflective enhanced aluminum coating on the hypotenuse, Tower Optical) was mounted on a custom stainless-steel mount (using UV-curable adhesive, Norland). This assembly was then positioned by aligning the front face of the microprism parallel to the caudal surface of the MEC and aligning the top surface of the microprism perpendicular to the (eventual) axis of excitation light propagation. A thin layer of Kwik-Sil was applied to the caudal MEC surface before microprism implantation to fill the void between the brain and the front surface of the microprism. The microprism and mount were rigidly held in place and the craniotomy sealed by application of a thin layer of Metabond to all exposed sides of the microprism (except the top surface of the prism) and mount and on any exposed skull or brain. A titanium headplate (9.5 mm × 38 mm) was then attached to the dorsal surface of the skull, centered upon and aligned parallel to the top face of the microprism. A titanium ring (27-mm outer diameter and 12.5-mm inner diameter, with a 3-mm high edge) was then attached to the top surface of the headplate, centered around the microprism, and the area between the craniotomy and the inner edge of the metal ring was covered with opaque dental cement (Metabond, Parkell, made opaque by adding 0.5 g of carbon powder, Sigma Aldrich).

#### Experimental Setup for DREADD inactivation experiments

Mice were head-fixed over a cylindrical treadmill (60 cm circumference and 10 cm width), which was enclosed in a box (60 cm length x 60 cm width x 63.5 cm height). After being head-fixed, an odor nozzle and lick spout were placed near the mouse. Odorized air was delivered using a flow dilution olfactometer^25^. The olfactometer consisted of two streams of air: a carrier stream (0.9 L/min) and an odorized stream (50 mL/min) which carried isoamyl acetate (2% isoamyl acetate in mineral oil; odorant from Cole-Parmer, 99+%). The two streams combined, and a solenoid valve was used to direct the odorized airflow either to the mouse (via the odor nozzle) or to a vacuum (1.8 L/min). Odor delivery was validated using a photoionization detector (PID). Licking was monitored throughout each training session and was detected using a capacitance sensor (SparkFun Capacitive Touch - AT42QT1010) with an electrode positioned on the lick spout. A solenoid valve was used to deliver water (∼6ul per drop) via the lick spout when appropriate. All experimental paradigms were automated and controlled using an Arduino Uno, and data collection was performed using a Picoscope oscilloscope (PICO4824, Pico Technology, v6.13.2) sampling at 1kHz. Mice were free to run on the treadmill during all training sessions. All training was performed in the dark, during the dark phase of the animals’ light cycle.

#### Behavioral Training

After recovering from surgery, mice began water restriction (∼1mL of water per day). Once mice reached ∼85% of their initial weight, they began pre-training for the tDNMS task. Mice were first acclimated to the experimental setup though a habituation phase. During habituation, series of 50 drops of water (3s apart) were delivered to the mouse (3-6 series of 50 drops per session). Habituation ended after mice licked to consume ≥80% of water drops in a series of 50 drops. Following habituation, mice began 3 phases of shaping: Shaping 1, Shaping 2, and Shaping 3. Shaping followed the same trial structure of the tDNMS task; however, only nonmatch trials were used. Each trial consisted of a flash of green light (lasting 0.25s in duration and preceding odor onset by 3s) to alert mice the trial was about to begin, the first odor, an interstimulus interval, the second odor, and a response window. Trials were separated by a random intertrial interval (ranging from 16-24s). In Shaping 1, a drop of water was automatically delivered 0.25s after second odor offset in each trial. Once mice licked to consume drops in ≥80% of trials, they progressed to Shaping 2. Probe trials were introduced in Shaping 2. During probe trials, the mouse had to lick within a 3s response window following second odor offset to trigger a reward. If the mouse successfully triggered reward, the next trial was another probe trial. If the mouse failed to trigger a reward in a probe trial, the following trial was automatically rewarded, after which the mouse was given another probe trial. Training on this phase continued until mice licked to earn reward in ≥20 consecutive trials. Mice then began Shaping 3, which had the same probe trial format as Shaping 2. However, in addition to licking in the response window, mice were also required to withhold licking during the first odor and interstimulus interval (ISI) to trigger reward delivery. Training on this phase continued until mice reached 2 consecutive sessions of ≥ 20 consecutive rewarded trials, after which they began the tDNMS task. Some mice failed to reach this benchmark yet routinely performed above chance on probe trials. These mice instead began the tDNMS task after reaching ≥ 80% correct performance on probe trials. Following shaping, mice began the tDNMS task, where match trials were introduced. Match and non-match trials were included in a pseudo-random manner and balanced so that half of trials were match and half were nonmatch. Nonmatch trials were evenly split between short-long and long-short trials. Mice were rewarded in nonmatch trials only if they withheld licking during the first odor and ISI and if they licked within the 3s response window following second odor offset. Mice received no reward in match trials and were punished with an increased ITI (+12s) for licking in the response window. Mice were trained for 1 session per day, with each session consisting of 100 trials for shaping, or 90 trials for the tDNMS task. Mice were trained at least 5 days/week during pretraining and 7 days/week during the tDNMS task.

Several cohorts of mice were used to establish the tDNMS task, with minor variations between cohorts. For the first cohort, the light cue indicating trial start was 0.25s before first odor onset, and each upcoming trial was assigned based on probability (25% chance of S-L, 25% chance of L-S, 50% chance of S-S). Subsequent mice were trained on the standard version of the task. In the standard task, the light cue indicating trial start appears 3s before first odor onset. Additionally, trials are presented in a block structure: each block of 4 trials consists of two S-S trials, one S-L trial, and one L-S trial, presented in a random order.

MEC inactivation experiments were performed in two cohorts of mice. Each cohort contained a balanced number of control and DREADD mice, with the experimenter blind to experimental condition. Following tDNMS training, the first cohort of mice was trained for an additional session of 30 trials with no odor (mineral oil only) to ensure mice use odor to solve the task. The second cohort of mice was instead trained on a Fixed Interval (FI) task^32^ following tDNMS training to determine if MEC is necessary to learn more rigid timing behavior. The same experimental apparatus was used for the FI task with the exception of odor delivery. In each session of the FI task, a (∼6ul) drop of water was delivered to the head-fixed mouse every 10s (150 drops per session). Mice were trained for 1 session per day for 5 days on the FI task. Schematics depicting tDNMS and FI behavioral paradigms (Fig. 1a, 4a, S1a, S12a&c) were made using BioRender.

Imaging data was collected from separate cohorts of mice. Imaging was performed using transgenic mice expressing GCaMP6s under the CaMKIIa promoter. Two mice were used to validate the efficacy of DREADD-mediated inhibition in MEC and were not trained on the tDNMS paradigm, while all other mice underwent tDNMS training. To determine how time cells adapted to changes in trial structure, mice were tested on probe sessions following tDNMS learning in which the ISI was lengthened from 2s to 5s on a subset of trials.

A final cohort of mice was trained on a version of the tDNMS task with modified durations. Due to the three trial type design, mice could solve the tDNMS task by learning to lick if total trial duration is long and to withhold if total trial duration is short. Standard experiments were performed with a 2s short odor, 3s ISI, and 5s long odor, making nonmatch trials 10s and match 7s. To test if mice used a rigid strategy, we decided to introduce probe trials that cause match and nonmatch trials to be the same overall duration. With our standard durations, this would mean lengthening the ISI on match probe trials by 3s. However, this change would increase task difficulty (increasing the time the mouse must resist impulsivity & increasing working memory demand), making it difficult to determine if a potential drop in performance is caused by increased task difficulty or use of a rigid strategy. To avoid this problem, we instead trained a separate cohort of mice on a version of the task with 3s short odors, 5s ISI, and 6s long odors so that we could reduce the ISI in nonmatch probe trials to equal total trial duration of match trials.

#### Two-photon imaging of MEC neurons

After mice were pretrained on the tDNMS task (14.5 ± 3.9 days of pretraining), we began two-photon laser resonance scanning of populations of neurons expressing GCaMP6s through the microprism using a Neurolabware microscope. Data were acquired with an 8 kHz resonant scanner, images were collected at a frame rate of 30 Hz with bidirectional scanning, and Scanbox software was used for microscope control and data acquisition. A Ti:Sapphire laser (Discovery with TPC, Coherent) at 920 nm was used as the excitation source, with average power measured at the sample (after the objective; 20×/0.45 NA air immersion objective (LUCPanFL, Olympus) with correction collar set at 1.25) of 50-120 mW. Imaging was also tracked using a PicoScope Oscilloscope (PICO4824, Pico Technology, v6.13.2) sampled at 1kHZ to synchronize data with behavior.

#### Histology

Following behavioral experiments, mice were perfused using 4% PFA in 0.1M PBS. The brain was removed and fixed in 4% PFA in 0.1M PBS for ∼24 hours. Brains were then rinsed 3x with 0.1M PBS, then stored in PBS for 1+ days before the tissue was sectioned in 50-100 micron sagittal slices using a vibrating microtome. Free floating slices were then incubated in 0.1M PBS with 0.1% Triton-X for 15 minutes, washed with 0.1M PBS and incubated for 3 hours in a 25:1 solution of 0.1M PBS with 435/455 blue or 530/615 red fluorescent NeuroTrace Nissl stain (Invitrogen). Brain sections were imaged and stitched using a VS200 Virtual Slide fluorescence microscope (Olympus) with a 10x OFN26.5, NA 0.40 objective (Olympus).

### Data analysis

#### General

Imaging data was analyzed on a IBuyPower Intel Core with Windows10 and custom software written in Matlab (2018b). No statistical methods were used to predetermine sample sizes. Sample sizes were based on reliably measuring experimental parameters while remaining in compliance with ethical guidelines to minimize the number of animals used. Repeated Measures ANOVA, paired t-test, Wilcoxon rank sum test, two-sample Kolmogorov-Smirnov test, chi-squared test, Wilcoxon signed-rank tests were used to test for statistical significance when appropriate, and all statistical tests were two-sided unless stated otherwise. For tests assuming normality, data distributions were assumed to be normal, but this was not formally tested. Experimenters were blinded during data collection of MEC - DREADD inactivation experiments in the tDNMS task (experimenters were not blinded for fixed interval experiments). All data in the text and figures are labeled as mean ± s.e.m. unless stated as mean ± s.d.

#### Behavioral Performance

Performance on the tDNMS task was analyzed by determining the percent of trials in which mice behaved correctly. Correct nonmatch trials were defined as those in which mice withheld licking during the first odor and ISI and licked in the 3s response window following second odor offset to trigger reward. Correct match trials were defined as those in which mice withheld licking for the duration of the trial. Mice that met the criteria to advance beyond shaping and begin the tDNMS task were included in analysis. However, a subset of mice were removed from analysis due to: a failure to retain the task structure learned in shaping (performance >3 SD below the mean on tDNMS session 1, n =1 mouse), headplate falling off (n = 1 mouse), or lack of virus expression (n = 2 mice). Though most behavior is analyzed as % correct trials, licking behavior was further examined in some instances. Licking was analyzed by identifying lick events, defined as samples (sampling rate of 1 kHz) in which the capacitance sensor detected a signal. For tDNMS behavior, any licking following reward delivery was removed to focus on predictive and not consummatory licking. Lick events were then binned in 0.25s bins, then normalized to the maximum within the session. Lick events were also used to examine performance in the Fixed Interval (FI) task. Performance on the FI task was measured as the % of trials in which mice engaged in predictive licking, where predictive licking is defined as an increase in lick events (determined by a significantly positive slope) in the 5s preceding reward delivery. Precision in the FI task was estimated by determining time of peak licking activity of each mouse on each session. Licking data was binned in 0.25s bins, and the bin number with the maximum number of lick events was recorded for each of the 150 trials. These values were then averaged and converted to time to give an estimate of peak lick time across the session.

#### Image processing, ROI selection, and transient analysis

In vivo two-photon data sets were acquired during the tDNMS task (120,000 frames per session). Movies were first motion corrected using whole-frame cross-correlation, as described previously^26^, and the motion-corrected time-series was used for all subsequent analysis. Regions of interest (ROIs) were defined using Suite2P v0.10.1^54^. Significant Ca2^+^ dF/F transients were identified using previously described methods^20,26,55^.

#### Defining time cells in the tDNMS task

Cells exhibiting significant tuning at specific time points during trials were defined based on mutual information^56,57^. Before computing mutual information, Ca^2+^ signals were normalized in each trial by dividing their peak dF/F value to prevent that one or two large transients determine a cell’s activity pattern. Subsequently, dF/F values were averaged across correct trials, and mutual information for the averaged dF/F was calculated using the following equation for each trial type (e.g., short-short, short-long, long-short):

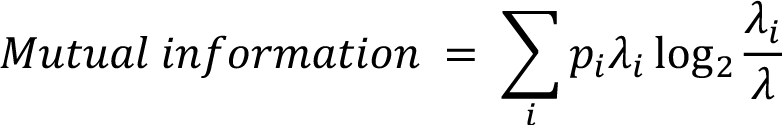

where *i* denotes bin, *p_i_* is occupancy rate in *i*th bin, *1_i_* is dF/F at *i*th bin, and *λ* is mean dF/F. To determine if the mutual information significantly exceeded that expected by random activity, the dF/F values for each trial were circularly shifted by a random amount, and mutual information was then computed using the shuffled data. Shuffling was repeated 1,000 times. The p-value was defined as the proportion of shuffled mutual information greater or equal to real mutual information. Cells were classified as active cells if mean dF/F > 0.03, and the active cells were classified as time cells if p-value of mutual information < 0.01.

#### Comparing correlation between match and nonmatch trial types across day 1 to day N

Time cells in either short-short or short-long trial types were selected, and the Pearson correlation coefficient between these two trial types was calculated for each cell. The same procedure was performed using time cells in either short-short and long-short trial types. All correlation coefficients were pooled and compared between Day 1 and Day N using Wilcoxon rank sum test.

#### Population vector differences between match and nonmatch trial types across trial epoch

Time cells in either short-short or short-long trial types were selected. Population vectors were constructed from dF/F of all selected cells at each time bin. The difference in dF/F between short-short and short-long trial types was then computed for each time bin. A corresponding shuffle distribution was generated by randomly assigning trial types and obtaining the dF/F difference 10,000 times. The time bin where the actual data exceeded the top 0.1% of the shuffled distribution was considered the significant discrimination timepoint for the trial type. The Discriminant Index was determined from the summation of the difference between the actual and averaged shuffle values from 8 to 11 seconds. This index represents the extent to which the actual dF/F distinguishes between trial types compared to the chance level. The same procedure was repeated for the time cells in either short-short or long-short trial types.

#### Decoding trial type using K-means clustering

K-means clustering was employed to assess the segregation of neural ensemble activity across different trial types. For each trial, population vectors were created by averaging dF/F during the late phase of trials (9∼11 seconds). To prevent overfitting, we reduced the dimensionality of the population vectors to 30 using Principal Component Analysis (PCA). The first two linear discriminant components were then determined, forming the axis for the two-dimensional Linear Discriminant Analysis (LDA) plot (Fig. 3g). Next, the K-means clustering method was applied to this two-dimension plot to classify trials into three clusters (Fig. S7). The decoding accuracy was assessed by calculating the proportion of correctly classified trials. To establish a baseline, the same procedure was repeated 10,000 times with shuffled trial types to generate a corresponding chance distribution. The p-value is then computed as the proportion of shuffle values equal to or greater than the actual accuracy of decoding.

#### Decoding trial type using Support Vector Machine (SVM)

A classifier for each Day N session was developed using Support Vector Machine (SVM) using the MATLAB function fitecoc. Only correct trials were used in this analysis. Mean dF/F of time cells during either the early phase of trials (0∼2 seconds) or the late phase of trials (9∼11 seconds) was used as an input matrix for the fitecoc function. The response input (Y input) for this function was either trial type (i.e., short-short, short-long, long-short) or match vs nonmatch of the training trials. The classifier computed from fitecoc then fed to the predict function to obtain decoded responses (e.g., short-short type) corresponding to the testing data set. We applied a leave-one-out cross-validation method, so the classification process was repeated as the number of entire trials. In each iteration, a single trial was selected for testing the data set and the rest of the trials were assigned for the training data set. To calculate the chance level of decoding accuracy shown in Fig. 3h, the response input for the classifier (i.e., trial types or match vs. nonmatch) corresponding to the training trials was shuffled when creating the classifier. This process was repeated 1,000 times to make a distribution of decoding accuracy for shuffled data. The bootstrap p-value was determined as the proportion of decoding accuracies in the shuffled distribution that were equal to or greater than the accuracy of the actual dataset.

#### Measuring activity coherence in error trials

Due to the low number of error trials in short-long and long-short trials, this analysis was only applied to short-short trial data. Correct trials were divided into two groups using a random subset of trials (Correct A and Correct B). The number of trials assigned to Correct B was set to match the number of error trials. Cell activity was sorted according to the sequence in Correct A and correlations for each cell’s activity were computed for Correct A versus Correct B (blue in Fig. 3j) and Correct A versus error (red in Fig. 3j). Random sampling of trials was repeated 1,000 times to obtain a distribution of correlation coefficients. Then, the mean values of these correlation coefficients were taken as the cell’s correlation coefficient values. To compare Day 1 with Day N, Pearson’s correlation coefficient between averaged dF/F on short-short error trials and correct trials of each trial type was calculated for short-short time cells (Fig. S8b). To ensure a consistent number of trials for comparison, seven trials were randomly selected and used to compute the averaged dF/F. The resulting sets of correlation coefficients were then compared across sessions (i.e., Day 1 vs Day N) and trial types using a two-way mixed ANOVA.

#### Comparing variance across time and distance

The dF/F values for time cells were re-charted based on the elapsed running distance from the moment the trial initiation light was turned on. For each trial, we measured the peak dF/F location within the distance dimension. The variability of these peak locations was assessed using the coefficient of variation, which is the ratio of the standard deviation to the mean and provides a standardized measure of dispersion. This method allows for the comparison of variations across different scales or dimensions. Following this, the procedure was applied similarly to measure the elapsed time since the initiation light was activated. The coefficients of variation for the time cells were then compared between the distance and time dimensions using a Student’s Paired T-test.

#### Generalized linear model

To determine the individual contributions of different behavioral variables (distance traveled (D), time elapsed (T), and licking (L)) in predicting the activity of MEC neurons, we employed a generalized linear model (GLM). This model fits calcium activity (dF/F) as a Gaussian linear function using various combinations of these three behavioral variables^29,30^. For the dF/F data of each cell in each trial type (S-S, S-L, & L-S), we developed seven models. These included three single-variable models (D, T, L), three double-variable models (TD, TL, DL), and one comprehensive model (TDL). We utilized data from 10 recording sessions to perform this modeling.

In our model, distance is represented by 10 binary variables, corresponding to 10 spatial bins, with each bin equal to one when the animal occupied that spatial bin and zero otherwise. Time and licking are represented in the same way by 10 and 2 binary variables, correspondingly. dF/F was then smoothed with a Gaussian kernel. Models were fit using the MATLAB *fitglm* function with a 5-fold cross-validation procedure. We then calculated the log-likelihood (LLH) increase for each model as follows: where N is the number of data points, 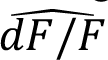 is the model predicted 𝑑𝐹/𝐹, and 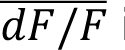 is the session mean dF/F. LLH is normalized by recording time (per min).

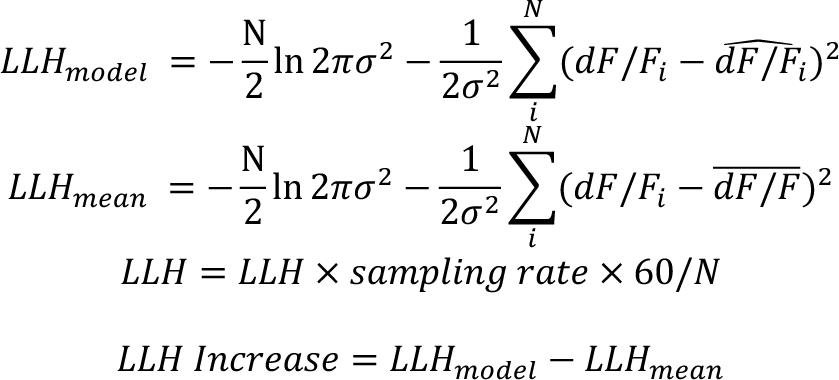

To select the model that best described the calcium activity of each cell, we first found the single variable model with best performance, then determined if any double variable models that included this single variable had better performance. If so, we then compared the double variable model to the full model. Model performance is determined by comparing the LLH increase using a one-sided signed rank test, with a significance value of p = 0.05. To ensure the significance of the models, we only use data from a cell if its best model has LLH increase > 0.

To estimate the specific contribution of each behavioral variable, we found all cells with significant full and double variable models. We then calculated the log-likelihood difference as follows (using time as an example):

If the best model of a cell is the full model:

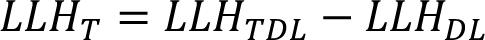

If the best model is one of the double variable models:

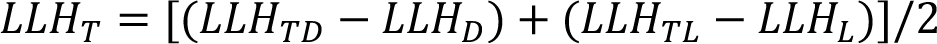

Across all cells and trial types, we identified N= 177 significant models (Kruskal-Wallis test followed by Wilcoxon rank-sum test with Bonferroni-correction, p<0.001 for all). We selected the model that best described the dF/F for each cell and trial type as the model with the highest LLH increase, which most often corresponded to a single-variable model of time (125 out of 177 significant models).

#### Poisson modeling of animals’ licking behavior

A Poisson GLM was fit to the anticipatory licking data individually for each mouse to assess animals’ internal model for solving the task. Behavioral data was down sampled to 10Hz and cropped to 20s per trial. The expected value for anticipatory lick counts, λ(t), was predicted according to a Poisson distribution:

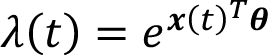

Where x(t) corresponds to the column vector of predictor features in the model at time t and 8 is the vector of weights assigned to each feature’s influence on the predicted lick count. Only anticipatory licks were used and any licks after water delivery were omitted. A no-strategy model was developed based on individual events-based cues, which could result in licking at key times throughout the trials but could never solve the task with 100% success. The 4 cognitive strategies (as outlined in the Results section) were tested by adding features that corresponded to whether the conditions given by an individual strategy were met. Each of these features were tested individually by adding them to the cue-based model and calculating the degree of improvement in the model fit to observed licking. For all models, cue-based features were lagged every time bin from 0 to 2 seconds and strategy-based features were given lags from 0 to 12 s to account for any licking between the offset of the second odor until the end of the trial. Estimated weights (𝜃^G^) for each feature were learned using python’s scikit-learn package. We applied L2 regularization to avoid overfitting due to predictor collinearity, and we evaluated the models by K-fold cross validation across trials (k=10). Models were scored using the deviance explained (d^2^) metric to assess goodness-of-fit. For models including strategy-based features, we calculated the Log-Likelihood increase (LLHi) over the cue-based model.

#### Comparing pairwise activity correlation between trial period and ITI

To measure the coherence of MEC time cells across tDNMS trial and non-trial epochs, we compared the pairwise activity of all time cell pairs between the trial period and the inter-trial interval (ITI)^58^. The entire time series of dF/F in a session (e.g., 120,000 frames) was segmented into 500ms time bins, and dF/F values were summed within each time bin. Time bins from 1 second before the first odor onset to the second odor offset (11 seconds) were included in trial period, and time bins from 5 seconds after the second odor offset to 4 seconds before the next first odor onset were included in ITI. Given the 3-5s gaps between each trial period and ITI, the likelihood of activity from one epoch influencing the other was minimal. The time bins within these gaps were excluded from the analysis. Kendall’s correlation (τ values) was calculated for the series of summed dF/F values during either the trial period or ITI across all pairs of simultaneously recorded time cells. Then, the coherence across trial and ITI epochs was measured by computing Pearson’s correlation coefficient between sets of corresponding τ values of the trial period and ITI (Fig. 6b). To generate shuffled data, the dF/F values were circularly shifted by a random amount before computing τ values. The same procedure was repeated for correct vs error trials (Fig. S15a) and between different trial types (Fig. S15b). In Fig. 6c, the τ values for the ITI are plotted against the difference in peak times of each cell pair during the trial period.

### Data availability

Data and code presented in this study are available upon reasonable request to the corresponding author and will be made publicly available 1 year after publication.

## Acknowledgments

We thank Daniel Dombeck, Michael Long, and Mark Sheffield for valuable comments on earlier versions of this manuscript. We thank Matt Wachowiak for generous support in designing and validating the olfactometers used in this study. This work was supported by the Whitehall Foundation, Brain and Behavior Research Foundation, NIH/NIMH 1 DP2 MH129958-01, NSF CAREER Award: IOS-2145814, and the University of Utah.

## Contributions

E.R.B, H.W.L and J.G.H. designed experiments and wrote the manuscript. E.R.B, H.W.L and J.S. collected data. E.R.B, H.W.L, J.S., J.C.B., and J.G.H. analyzed and interpreted data. E.R.B. and J.G.H. built equipment used to collect data.

---

**Figure S1.**
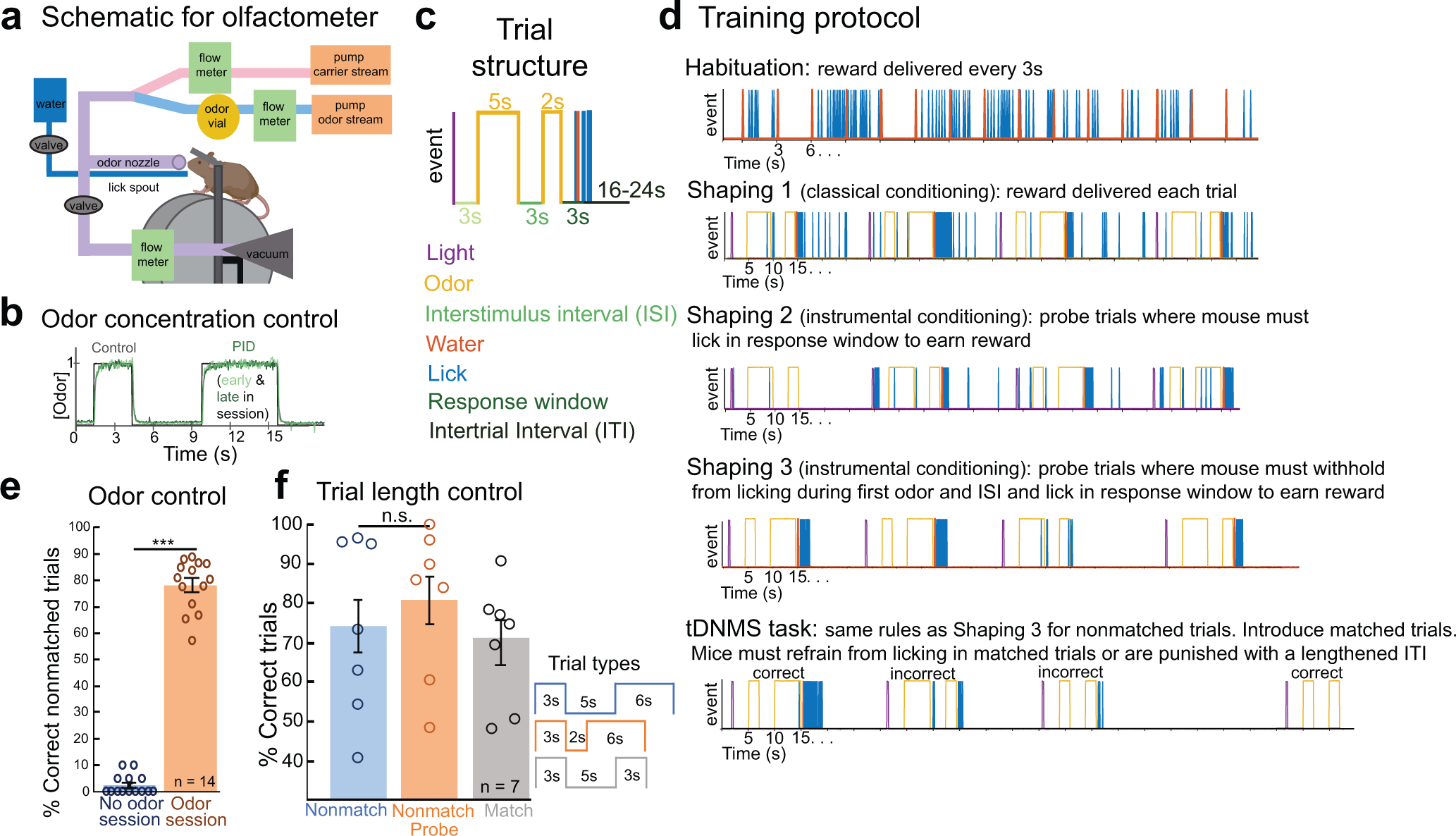
tDNMS set-up and controls. **a.** Experimental set-up. Odorized air is directed either to the mouse or to a vacuum. A lick spout, connected to a capacitance sensor, delivers water and is used to monitor mouse licking. **b.** Odor concentration control. Odor concentration was measured using a photoionization detector (PID). Odor can be delivered with high temporal specificity at a constant concentration over 45 minutes, as shown by PID measurements (green) relative to control signal (black). **c.** Trial structure. Each trial consists of two presentations of the same odor, each for either a 2 or 5s duration, separated by an interstimulus interval (ISI). Trial start is signified by a visual cue, and trials are separated by a 16-24s intertrial interval. **d.** Training protocol. Mice undergo three phases of pretraining (see Methods). **e.** Odor control session. After completing tDNMS training, mice were tested with no odorant (mineral oil only). Mice failed to solve nonmatch trials in the absence of odor but performed well in a prior session in which odor was used (p<0.001, paired t-test; n=14 mice). Bars represent mean ± s.e.m.. **f.** Trial length control session. A cohort of mice was trained on a version of the tDNMS task with modified durations. The ISI was then manipulated on a random subset of nonmatch trials (“probe trials”) so that overall trial duration was identical to match trial duration. If mice use total trial duration to solve the task, rather than individual stimulus durations, they should incorrectly withhold licking on probe trials. Instead, there was no significant difference between standard nonmatch and probe trial performance (p=0.19, paired t-test; n=7 mice). Bars indicate mean ± s.e.m.

**Figure S2.**
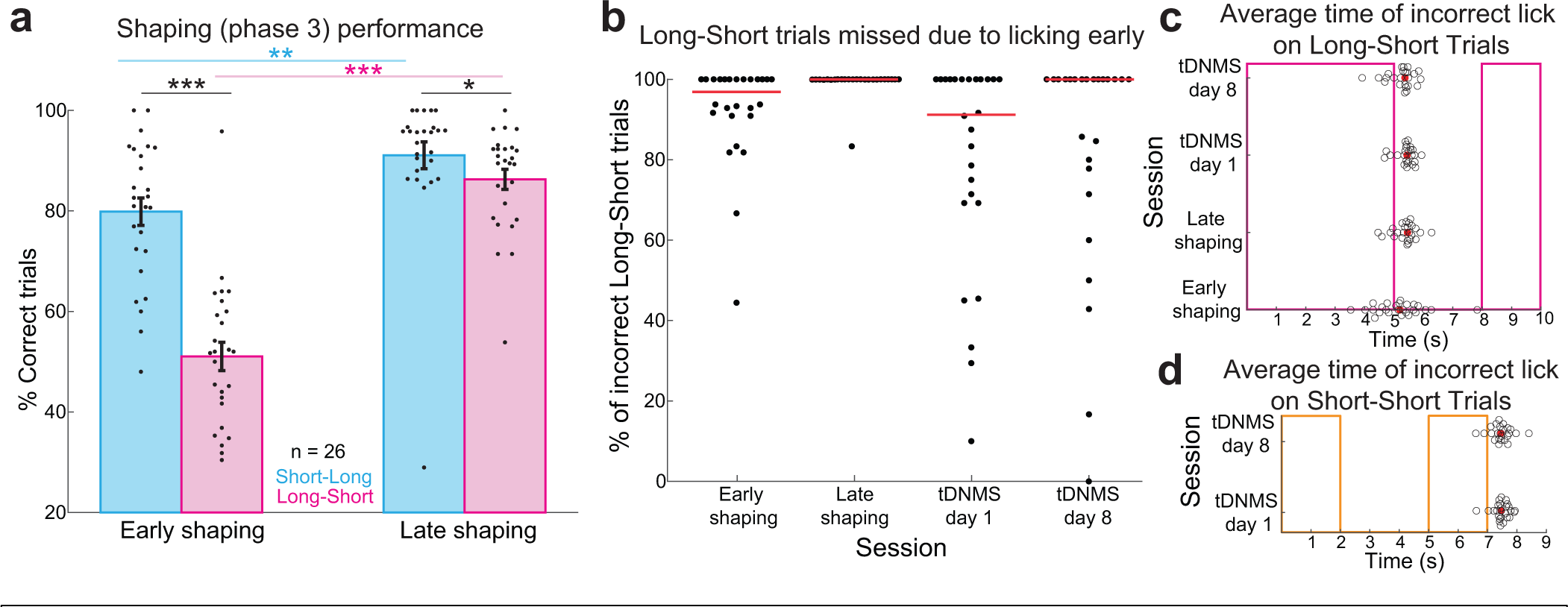
Additional behavioral analysis of mice in Figure 1. **a.** Average performance by trial type during shaping (phase 3). Shaping consists of probe trials where mice must correctly trigger reward and automatic trials where reward is automatically delivered. Performance was examined on all probe trials within the first 1/2 session of shaping phase 3, termed “early shaping”, and the last 1/2 session of shaping phase 3, or “late shaping”, for each mouse. Dots represent performance of each mouse, and bars show mean ± s.e.m. across mice. Mice performed better on short-long trials than long-short both early (p<0.001, paired t-test; n=26 mice) and late (p=0.04, paired t-test) in shaping. Additionally, performance was higher for both short-long (p<0.01, paired t-test) and long-short (p<0.001, paired t-test) trials in late compared to early shaping. **b.** Reason for mistakes on long-short trials. During shaping phase 3 and the tDNMS task, mice can miss nonmatch trials either by withholding licking or by licking prematurely during the first odor and/or interstimulus interval. The percent of incorrect long-short trials in shaping phase 3 and the tDNMS task missed due to licking early is shown. Dots indicate values for each mouse, with red lines showing the median value across mice. **c.** Average time of first incorrect lick on long-short trials relative to first odor onset during shaping phase 3 and tDNMS task. Black dots represent the average time of first lick across all incorrect long-short trials for a given mouse, and red dots show the median value across mice. **d.** Average time of first incorrect lick on short-short trials in the tDNMS task relative to first odor onset. Black dots represent the average time of first lick across all incorrect short-short trials for a given mouse, and red dots show the median value across mice.

**Figure S3.**
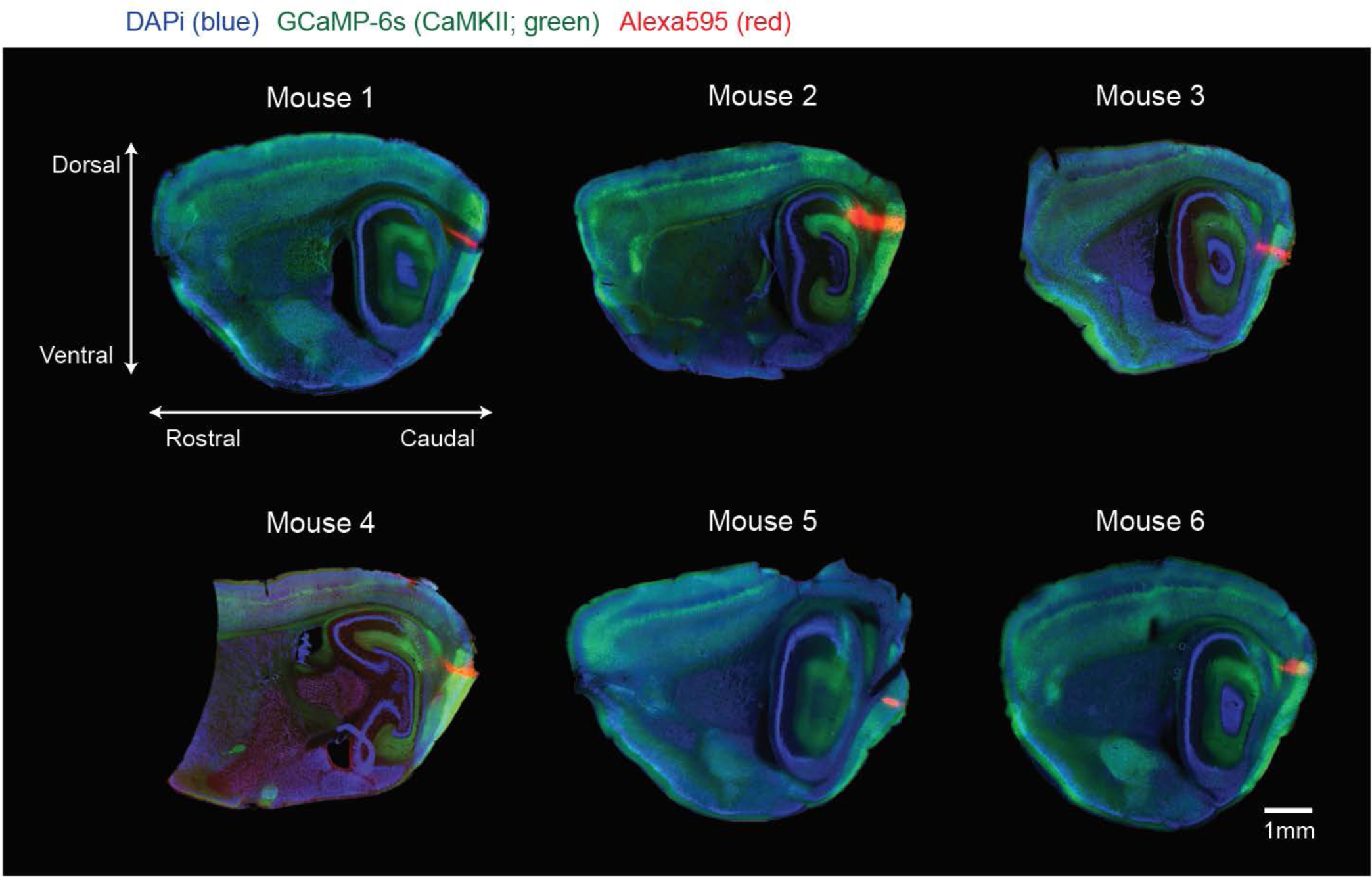
Histological verification of in vivo imaging in the MEC. Sagittal sections of post-mortem histology from all six mice utilized in the in vivo calcium imaging experiments. MEC neurons labelled with GCaMP-6s (green). The sections are stained with NeuroTrace 435/455, displaying neuronal morphology in blue. The approximate locations of the two-photon imaging fields of view (FOV) for each mouse are labeled with red Alexa594. This labeling was achieved by inserting a pin coated with Alexa594 at sites corresponding to prominent vascular landmarks visible both in the in vivo two-photon imaging and under a dissecting scope during the ex vivo marking procedure. Confirmation of the imaging sites within the MEC was based on the presence of the lamina dissecans, the relative position of the post-rhinal border to the pin mark, and the characteristic circular shape of the dentate gyrus as observed in the medial-lateral sagittal sections.

**Figure S4.**
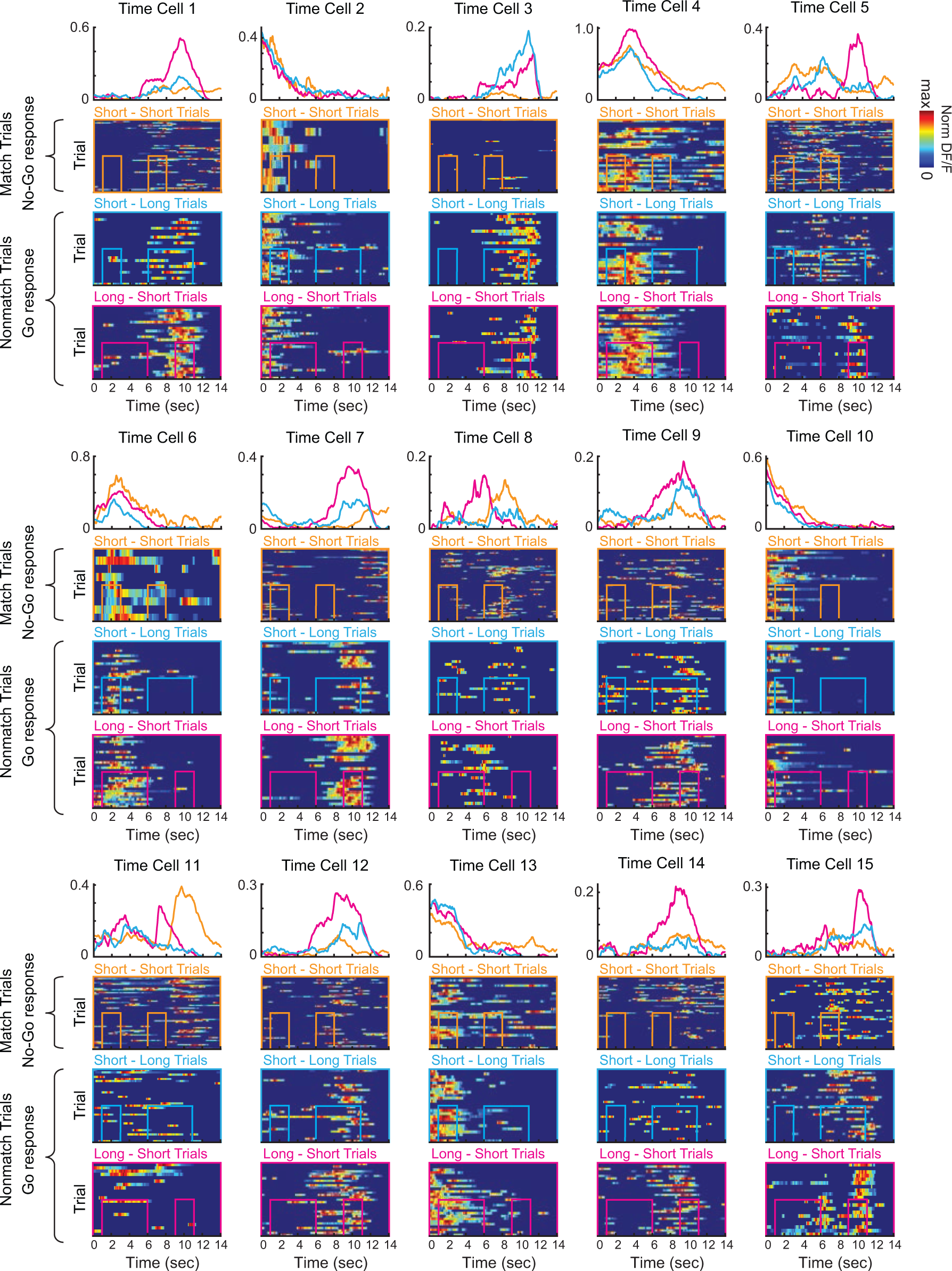
Additional examples of MEC time cells. For each time cell, mean dF/F displayed for each trial type (top) and dF/F activity on each trial, sorted by trial condition (below).

**Figure S5.**
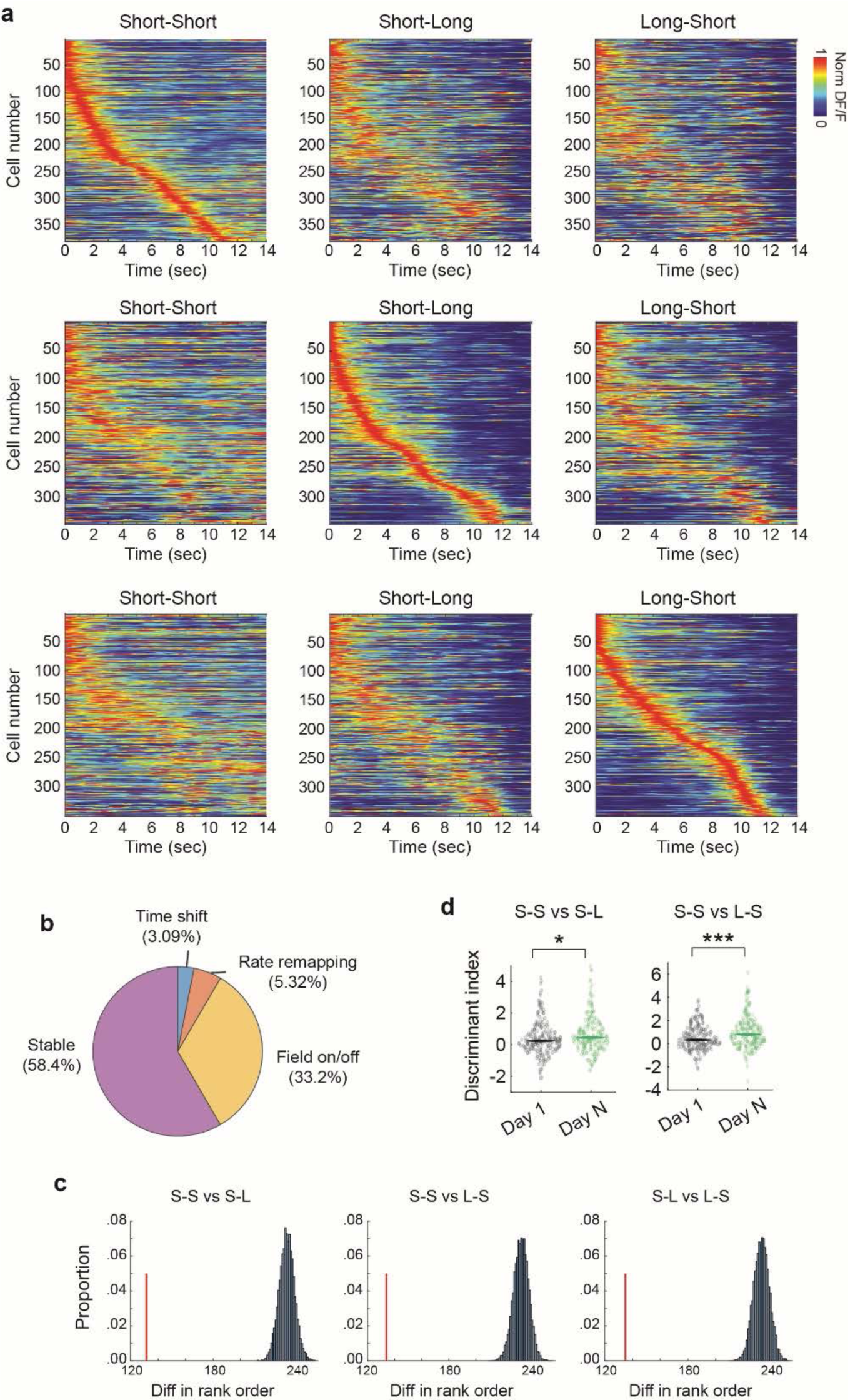
All MEC time cells sorted for each trial type. **a.** Top, sequence of MEC time cells significantly tuned for SS trials, sorted by SS trials and displayed for SS, SL and LS trials. Middle, same as above, expect for significant MEC time cells on SL trials and sorted by SL trials. Bottom, same as top, except for significant MEC time cells on LS trials and sorted by LS trials. **b.** Proportion of MEC time cells that either remained stable, displayed a time shift, displayed rate remapping, or displayed on/off dynamics across trial types. **c.** Rank order analysis for shuffle distribution (black) and real data (red). The similarity of the sequences of time cells across trial types is examined by comparing their rank orders. Each time cell is assigned three rank orders, corresponding to its sorting by peak timing for each trial type. Subsequently, the mean difference between rank orders within a cell is compared to a shuffle distribution, generated by shuffling rank order of cells 10,000 times. The p-value is computed as the proportion of shuffle values smaller than the actual data. Notably, p-values are zero for all three comparisons. **d**. Discriminant Index indicates the extent to which the difference in dF/F between trial types deviates from chance level. S-S vs S-L: Day 1 n = 224, Day N n = 221, z = 2.55, p = 0.01; S-S vs L-S: Day 1 n = 225, Day N n = 231, z = 3.80, p < 0.001, Wilcoxon rank sum test. Individual data points with median.

**Figure S6.**
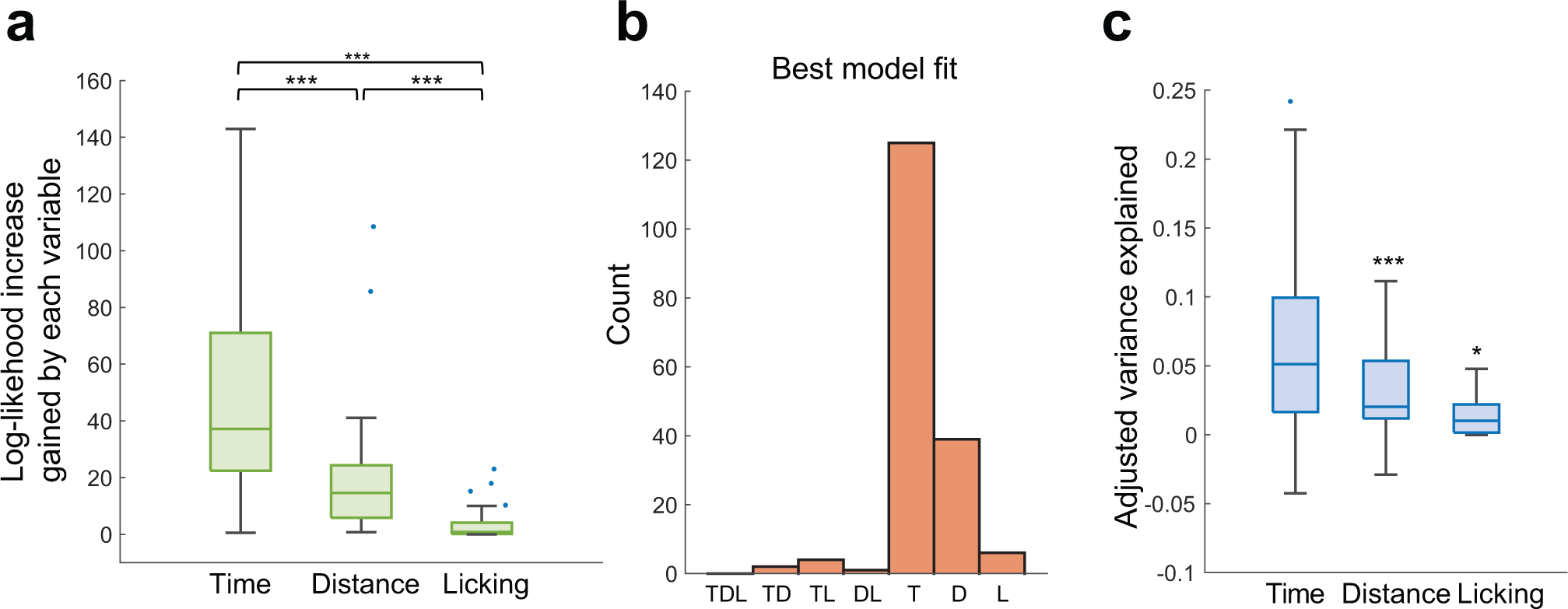
Generalized linear model demonstrates neurons are tuned to time. A generalized linear model was used to assess whether neurons are tuned to one of three variables-time in the trial, distance travelled from trial start, or licking – or a combination of 2 or 3 variables (see Methods). Analysis performed on n=695 time cells, collected from 10 behavioral sessions, lead to n=177 significant models. **a.** Boxplot showing loglikelihood increase gained by each variable: time (median = 37.15), distance (median = 14.63) and licking (median = 0.82) (Kruskal-Wallis test, p<0.001; followed by Wilcoxon rank-sum test with Bonferroni-correction: Time vs. Distance: p<0.001; Time vs. Licking: p<0.001; Distance vs. Licking: p<0.001). Log-likelihood was normalized to recording time in minutes. **b.** Histogram demonstrating the model that best described the calcium activity of each cell and trial type. **c.** Boxplot showing adjusted variance explained for models that best describe the calcium activity of each cell for the single variable models. Number of models n = 125,39,6. (Wilcoxon signed rank test (median greater than 0), p<0.001, p<0.001, p=0.03). Blue dots in **a** and **c** represent outliers.

**Figure S7.**
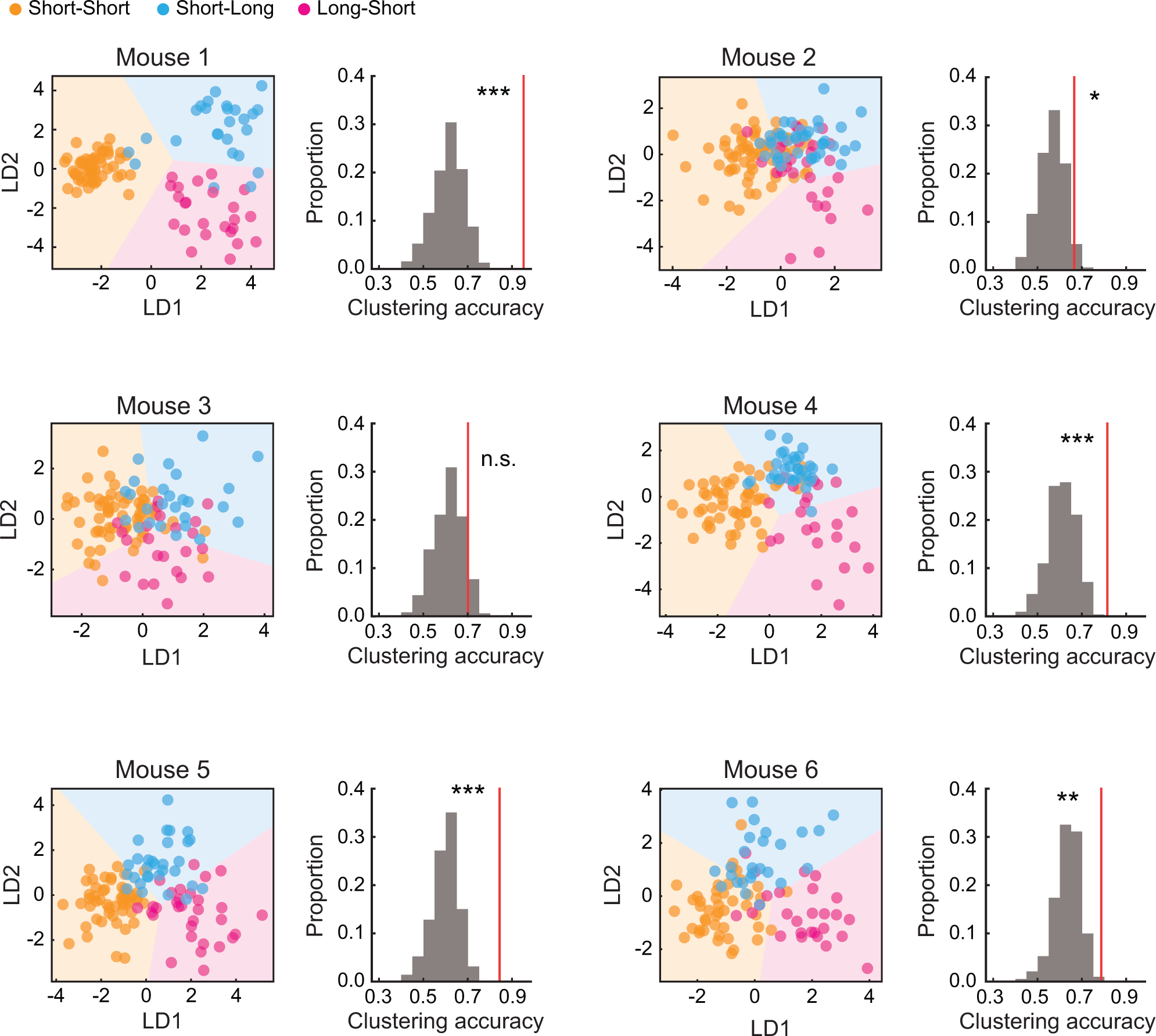
Trial type decoding analysis for each mouse. For each panel: Left, LDA plots for each mouse on each trial type (SS – orange, SL – Blue, LS – Magenta). K-means clustering is then applied to the LDA plots to categorize the dots into three clusters. The background colors indicate the clustering result. The accuracy of clustering analysis is determined by the proportion of dots correctly classified into their respective trial type. Right, clustering accuracy is compared between the bootstrapped shuffle distribution with randomly assigned trial labels (grey) and the actual data (red). The p-value is computed as the proportion of shuffle values larger than the actual data. For mouse 1 through 6: p = 0, 0.04, 0.08, 0, 0, 0.001, respectively.

**Figure S8.**
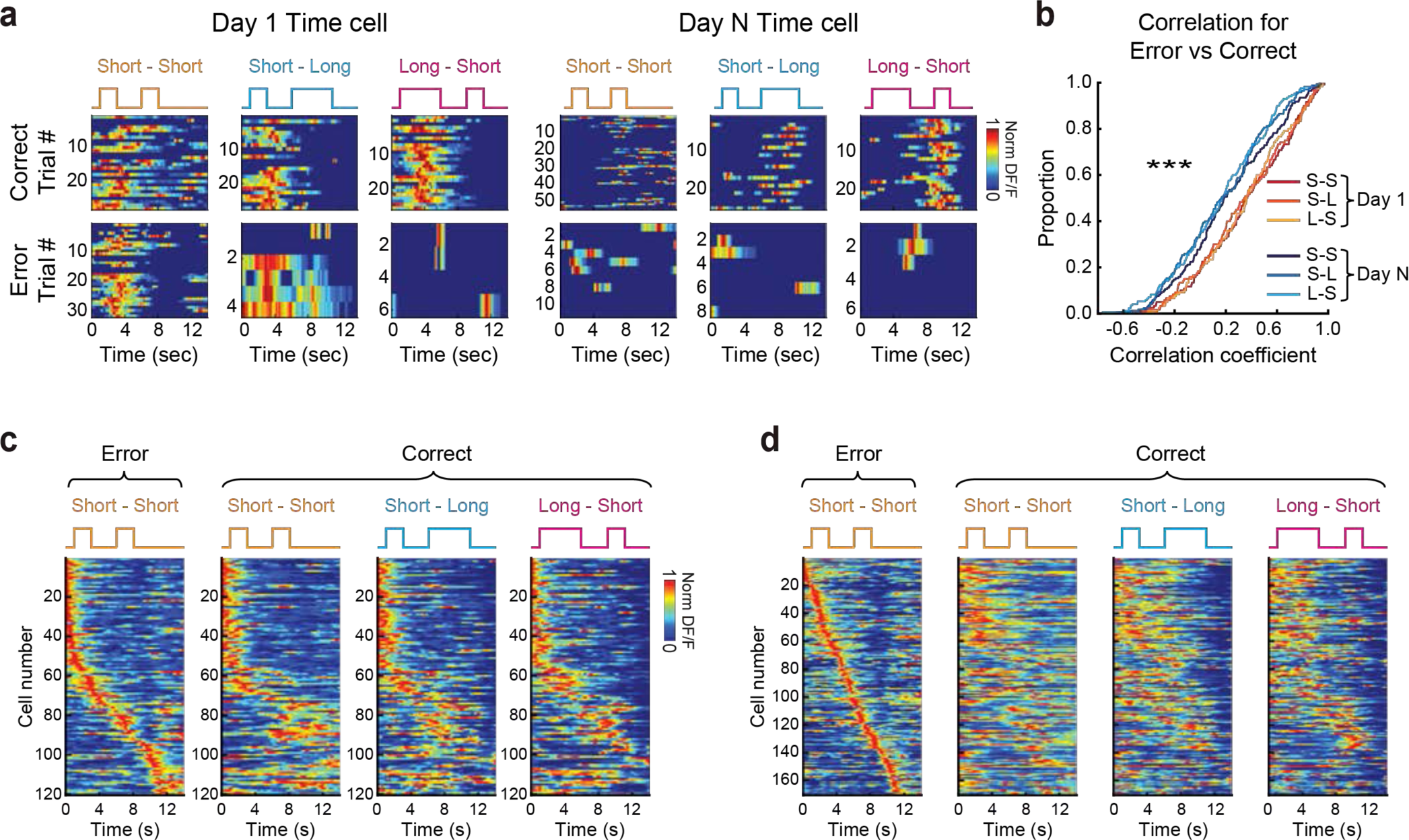
Analysis of MEC time cell activity on correct and error trials. **a.** Representative examples of MEC time cells on day 1 (left) and day N (right), depicting activity for both correct and error trials across all three types of trials. **b.** Cumulative distribution functions of correlation coefficients calculated for each MEC time cell, comparing activity across different trial conditions on day 1 and day N. session type main effect: p < 0.0001, F(1, 288) = 18.5; trial type main effect: p = 0.12, F(1.93, 555.15) = 2.1; interaction: p = 0.33, F(1.93, 555.15) = 1.1, two-way mixed ANOVA with trial type and session factors. **c.** Sequence of activity of MEC time cells recorded on day 1, arranged according to their activity during error trials on Short-Short (SS) trials. This sequence is then applied to display cell activity for all three trial conditions (SS, Short-Long (SL), and Long-Short (LS)) during correct trials, maintaining the order from the error trials. **d.** Same as in c, but for recordings on day N.

**Figure S9.**
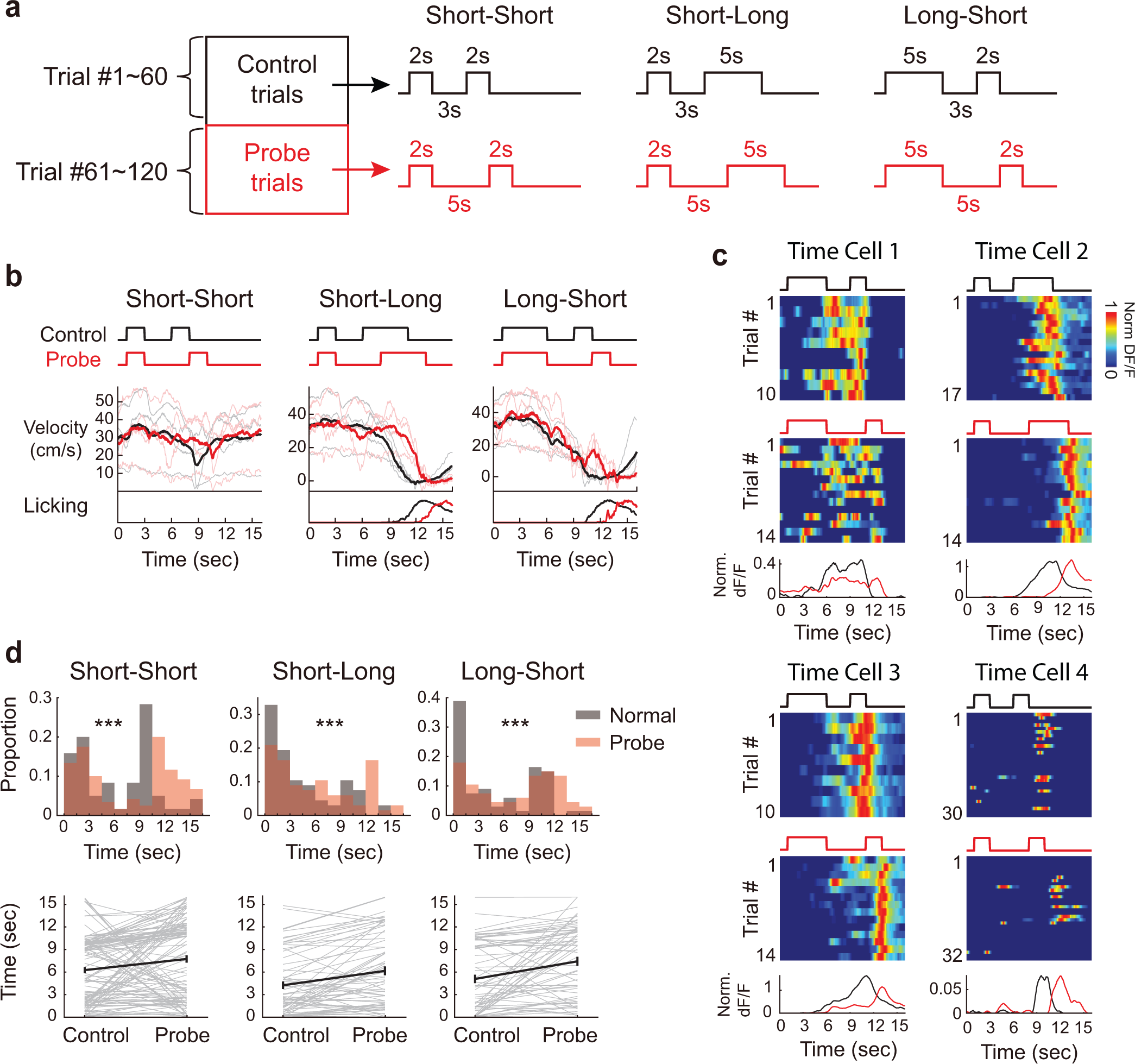
MEC time cells on ISI probe trials. **a.** Schematic for probe trials. **b.** Comparative analysis of mean velocity and licking behaviors under three different conditions: Short-Short, Short-Long, and Long-Short, during standard (black) and probe (red) trials. **c.** Four example MEC time cells during control (black) and probe (red) trials. **d.** Aggregated data showing the timing of peak responses across the MEC time cell population under Short-Short, Short-Long, and Long-Short conditions, compared between standard (black) and probe (red) trials. Short-Short trial types: n = 120, p < 0.0001 z = 3.9; Short-Long trial types: n = 67, p < 0.0001, z = 4.5; Long-Short trial types: n = 67, p < 0.0001, z = 4.2, Wilcoxon signed-rank.

**Figure S10.**
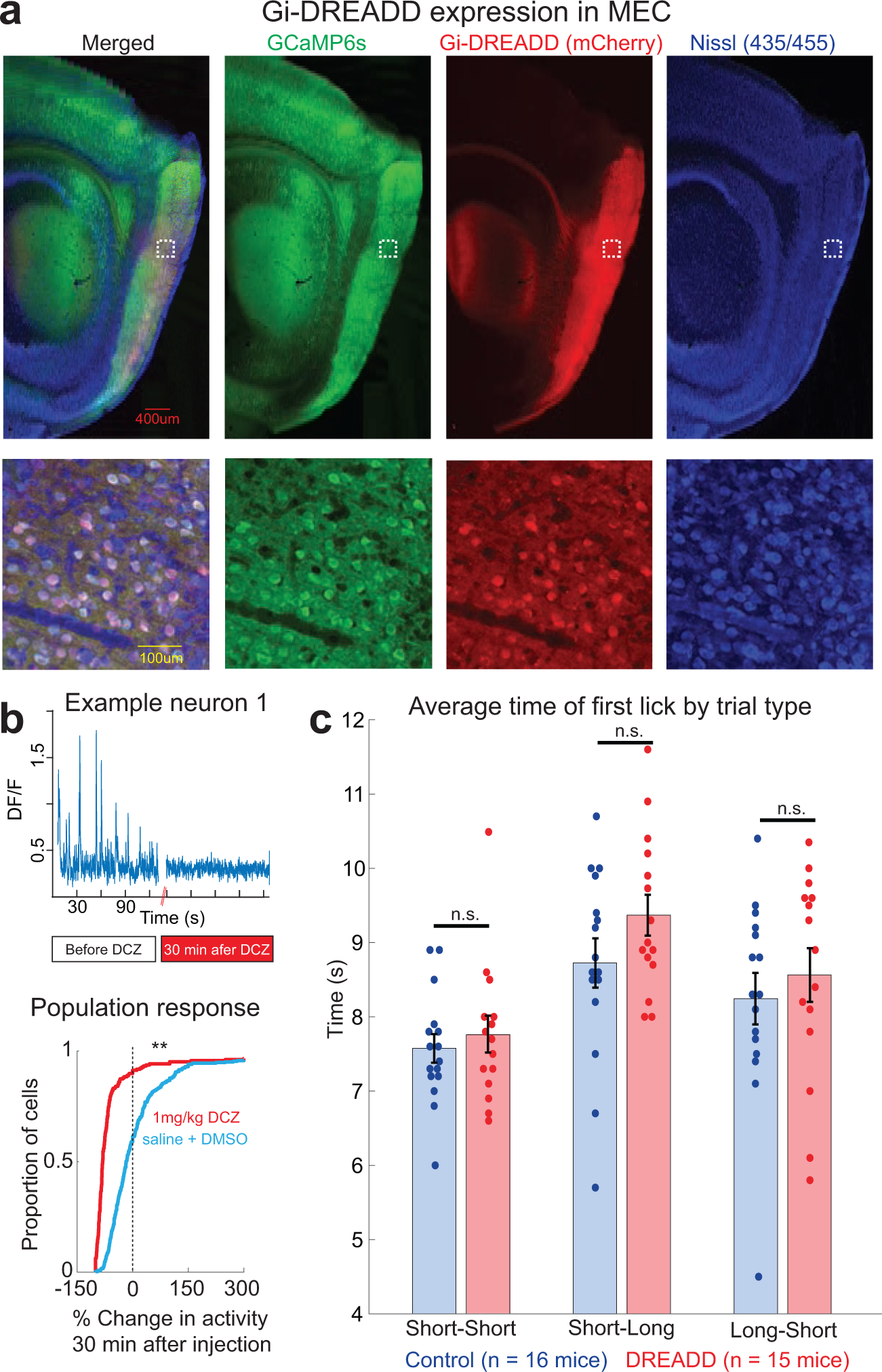
MEC DREADD inactivation. **a.** Histology showing co-expression of GCaMP6s and hM4D(Gi)-mCherry in MEC**. b.** Simultaneous in-vivo 2-photon GCaMP6s imaging and hM4Di inactivation in MEC. Activation of inhibitory DREADDs by 1 mg/kg I.P. injection of DCZ reduces average number of Ca2+ transients in MEC neurons by 80% at 30 minutes post injection compared to before DCZ injection. Left, example neuron before and after DCZ administration. Right, population response. In both the control (blue) and DCZ (red) conditions, GCaMP activity was monitored over 5 minute periods, and the change in activity was measured for each cell (n=302 neurons in 2 mice; p<0.01, Kolmogorov-Smirnov test). **c.** Average time of first lick relative to first odor onset on session 1 of the tDNMS task. Dots show average time of first lick for each mouse, with bars showing mean ± s.e.m. across mice. There is no difference in average time of first lick for DREADD (n = 15) and Control (n = 16) mice in any trial type (Short-Short: p=0.56, Short-Long: p=0.14, Long-Short: p=0.52, p values calculated using unpaired t-tests).

**Figure S11.**
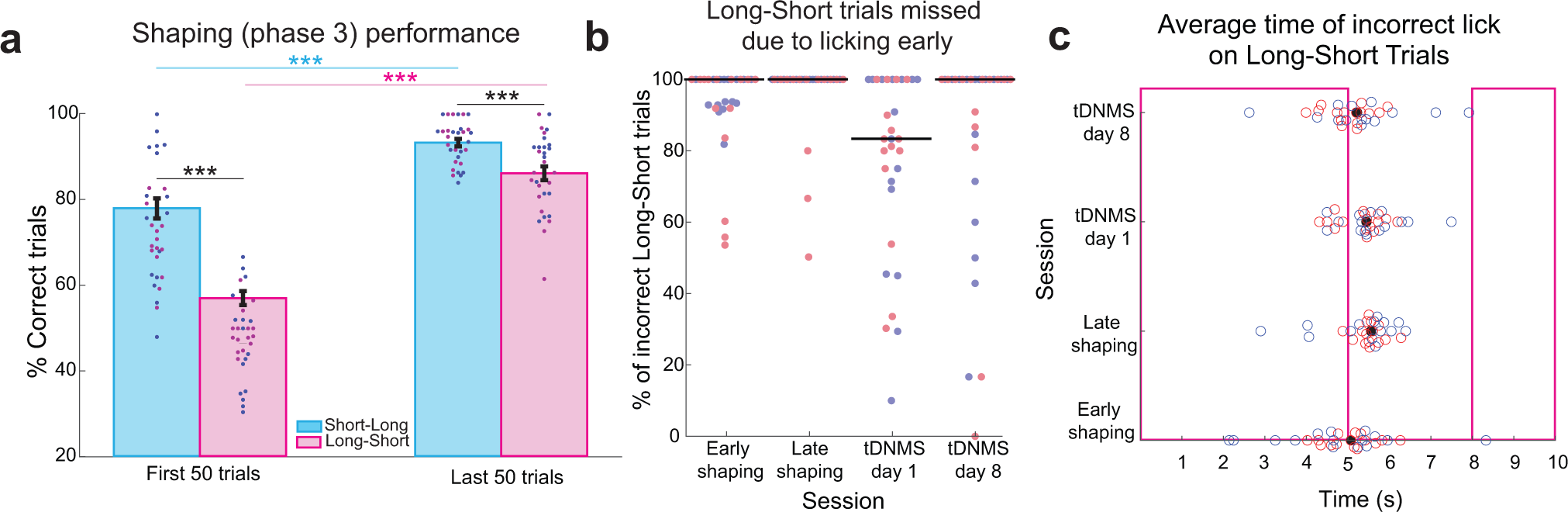
Additional behavioral analysis of mice in Figure 4. **a.** Average performance by trial type during shaping. Shaping consists of probe trials where mice must correctly trigger reward and automatic trials where reward is automatically given. Performance was examined on all probe trials within the first 1/2 session of shaping phase 3, termed “early shaping”, and the last 1/2 session of shaping phase 3, or “late shaping”, for each mouse. Mice performed better on short-long trials than long-short both early (p<0.001, paired t-test; n=31 mice) and late (p<0.001, paired t-test) in shaping. Additionally, performance was higher for both short-long (p<0.001, paired t-test) and long-short (p<0.001 paired t-test) trials in late compared to early shaping. Dots represent performance of each mouse, with blue dots for Control mice (n = 16) and red for DREADD mice (n = 15). Bars show mean ± s.e.m. across all mice. **b.** Reason for mistakes on long-short trials. During shaping phase 3 and the tDNMS task, mice can miss nonmatch trials either by withholding licking or by licking prematurely during the first odor and/or interstimulus interval. The percent of incorrect long-short trials in shaping phase 3 and the tDNMS task missed due to licking early is shown. Dots indicate values for each mouse, with DREADD mice shown in red and Control in blue, and black lines show the median value across all mice. **c.** Average time of first incorrect lick on long-short trials relative to first odor onset during shaping phase 3 and tDNMS task. Blue (Control) and red (DREADD) dots represent the average time of first lick on all incorrect long-short trials for a given mouse, and black dots show the median value across all mice.

**Figure S12.**
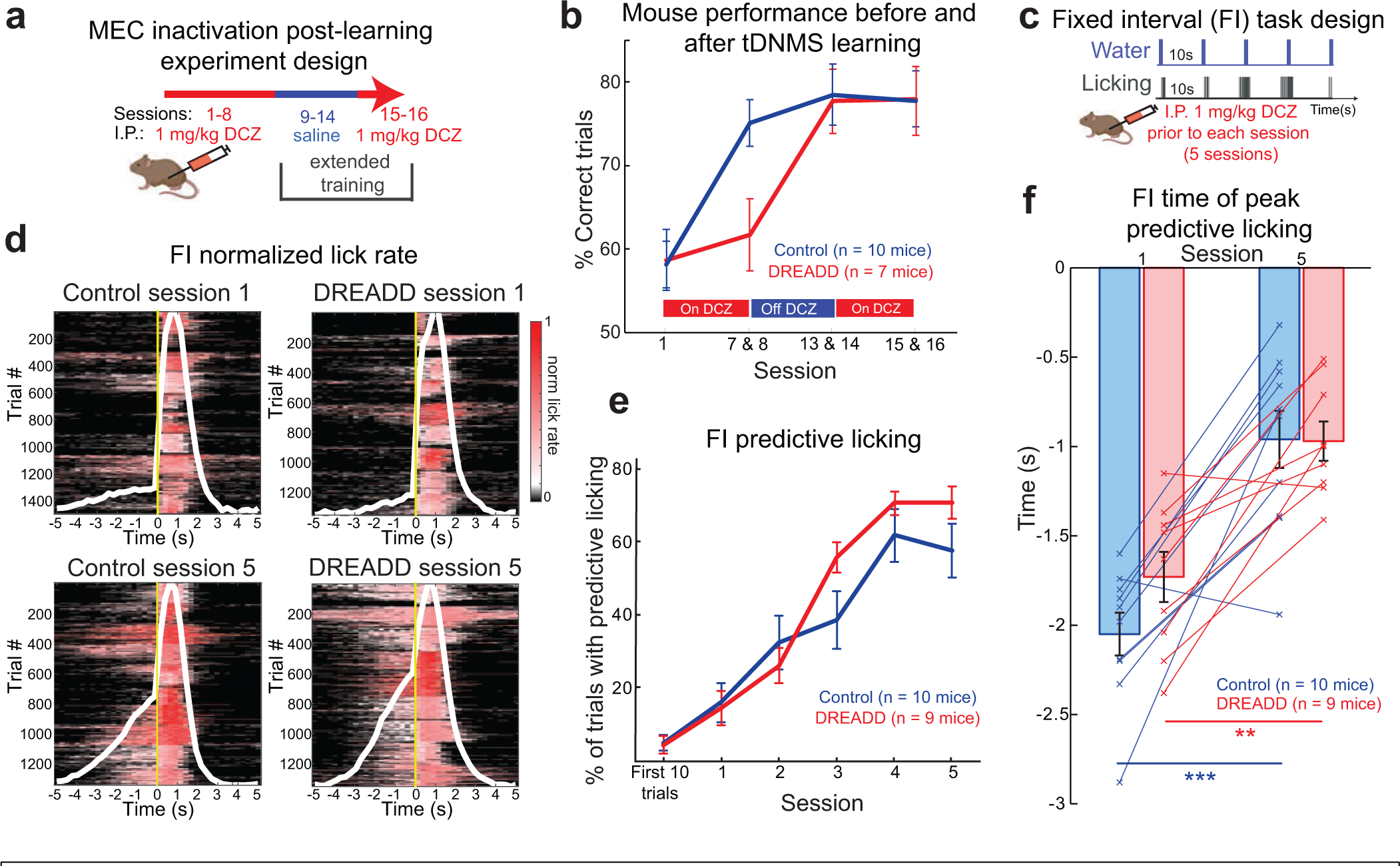
MEC is not required for all interval timing behavior. **a.** Schematic for inhibiting MEC after learning in the tDNMS task. After experiments testing the role of MEC during learning (Figure 4), a subset of mice underwent extended training to determine whether MEC is necessary for ongoing task performance. **b.** Though MEC inhibition impaired learning in the tDNMS task (Sessions 1-8), DREADD mice learned the task in the absence of MEC inhibition (Sessions 9-14). Following learning, subsequent administration of DCZ to inactivate MEC did not affect performance in Sessions 15-16. **c.** Fixed interval task schematic. MEC DREADD (n = 9) and Control (n = 10) mice were trained on a fixed interval (FI) task (Toda et al. 2017). A droplet of water (4-6ul) was delivered every 10s to head-fixed mice. Licking was measured; time-locked predictive licking indicates learning the timing of water delivery. The DREADD agonist DCZ (1 mg/kg) was delivered 5 min prior to each session. **d.** Licking behavior of DREADD (n=9) and Control (n=10) mice on sessions 1 and 5 of the FI task. Licking was normalized to the maximum lick frequency with each session for each mouse. All trials for all mice are shown; water delivery occurs at 0s, indicated by a yellow line. Average lick response for each session is shown (white). **e.** Fixed interval learning. Predictive licking is defined as an increase in lick rate, measured over 5 seconds preceding the upcoming reward delivery. Both DREADD and Control mice learn the temporal structure of the task, as demonstrated though more frequent engagement in predictive licking from sessions 1-5. Data represent mean ± s.e.m. across mice. **f.** Average time of peak licking activity in FI task. From session 1 to 5, peak licking activity moves closer to upcoming reward delivery for both Control (p < 0.001, paired t-test) and DREADD mice (p < 0.01 paired t-test). Data points represent average time of peaking licking on Session 1 and 5 for each mouse; bars indicate mean ± s.e.m. across mice.

**Figure S13.**
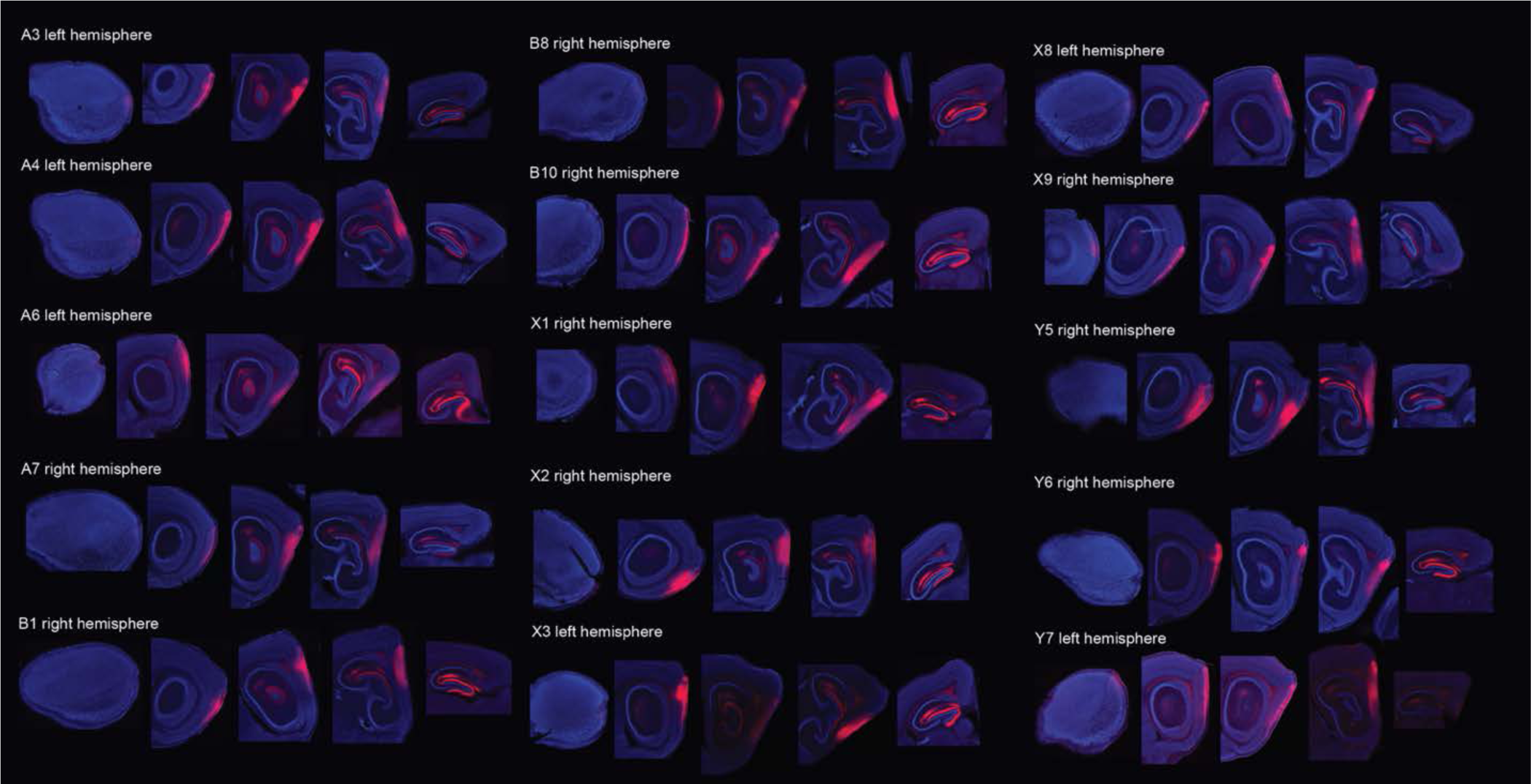
Histology showing expression of hM4D(Gi)-mCherry in MEC. Injections were performed bilaterally; one hemisphere is shown per mouse (mouse identity and hemisphere noted). Five sagittal sections are shown per mouse, ranging from lateral (left) to medial (right). The middle three sections include MEC.

**Figure S14.**
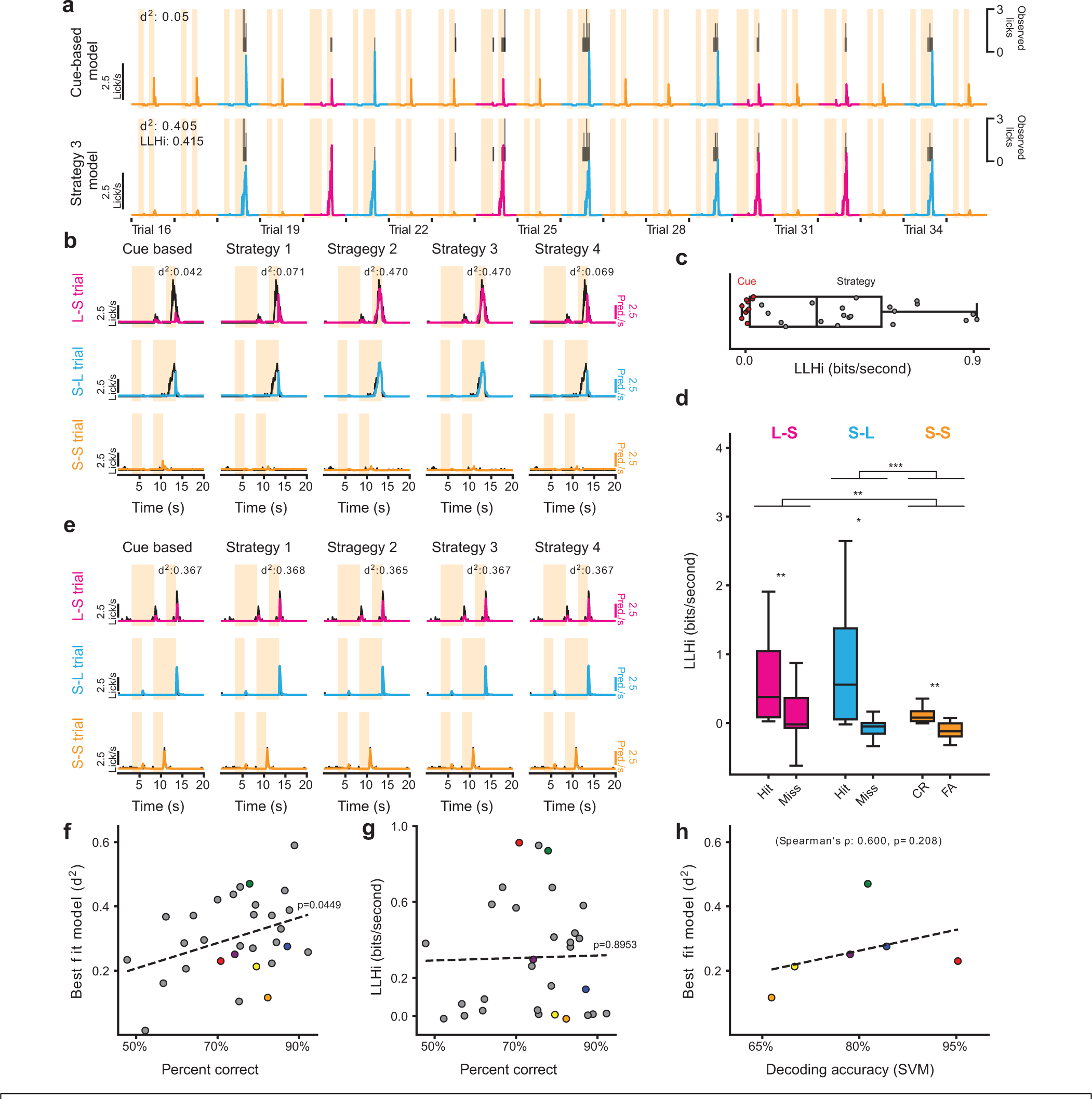
Additional details on Poisson Regression analysis of behavior. **a**. Example of modeling behavior during tDMNS task. *Top.* Baseline cue-based model including only odor offset, long cue, and second cue features. *Bottom.* Strategy-based model. Observed anticipatory licks are in black above each model’s predicted value. **b**. Comparison of cue-based and strategy-based models for an individual animal. Mean lick rate for each trial type in black, model predictions in color. **c**. Fitting a Gaussian Mixture Model with 2 components to the best fitting LLHi over the cue-based model reveals two populations: one cue-based where additional features do not improve model performance (n=10 mice) and one strategy-based with improved model predictions (n=20 mice). **d**. A strategy-based model fits better on hit/correct reject trials compared to miss/false alarm (based on Strategy 3) (Two-factor ANOVA significant effect for trial type F(2,155)=7.64, p <0.001 and result F(1,155)=35.6, p <0.001, but not type x result F(5,155) = 2.26, p=0.11, post-hoc tests: * - p<0.05, ** - p<0.01, *** - p<0.001). **e**. Example cue and strategy-based models for an animal where both fit similarly. **f**. Relationship between average mouse performance on the task and how well models fit the data (linear regression, r=0.369, p=0.045). **g**. No significant relationship between average percent correct and evidence for animals using a strategy-based solution, based on best fitting strategy (linear regression, r=0.025, p=0.895). **h**. No significant relationship between ability to decode trial type from the neural data and the ability to fit a model to the behavior, based on best fitting strategy (strategy and cue-based models, ρ=0.600, p=0.208, Spearman’s rank correlation coefficient). Colors in panels **f**-**h** indicate imaged mice, gray indicates not imaged.

**Figure S15.**
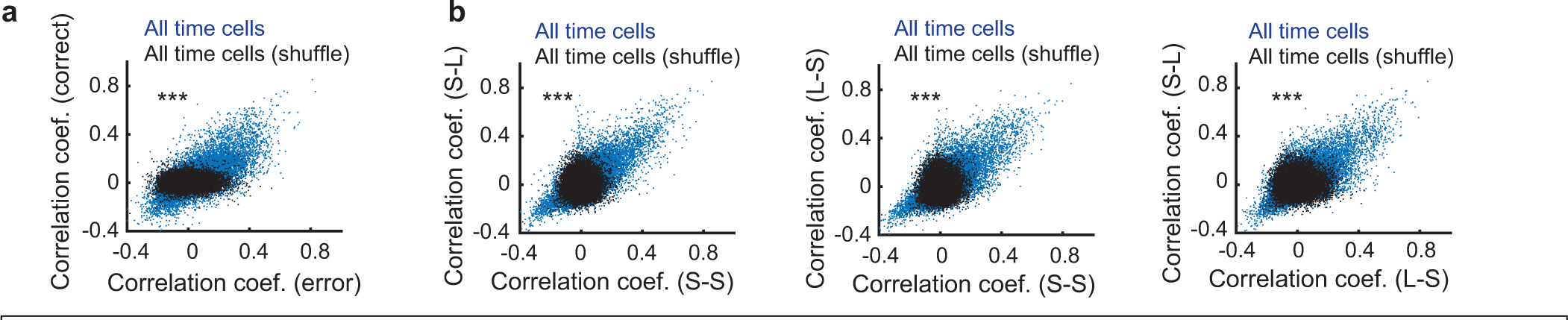
Activity correlation between time cells maintained across conditions. **a.** Pairwise correlation between all time cells during correct trials and error trials in the tDNMS task shown for real (blue) and shuffled data (black). n = 15,727, z = 69.6, p < 0.0001, Chi-squared test. **b.** Same as in a, but shown for correct trials on each trial type (S-S vs S-L, right; S-S vs L-S, middle; L-S vs S-L, right). S-S vs S-L: n = 15,727, z = 72.4, p < 0.0001; S-S vs L-S: n = 15,727, z = 66.3, p < 0.0001; L-S vs S-L: n = 15,727, z = 65.8, p < 0.0001, Chi-squared test.

## References

1. Tulving, E. Précis of Elements of episodic memory. Behav. Brain Sci. 7, 223–238 (1984).

2. Tulving, E. Episodic and semantic memory. in Organization of memory xiii, 423–xiii, 423 (Academic Press, 1972).

3. Kimble, D. P. The effects of bilateral hippocampal lesions in rats. J. Comp. Physiol. Psychol. 56, 273–283 (1963).

4. Niki, H. Response perseveration following the hippocampal ablation in the rat. Jpn. Psychol. Res. 8, 1–9 (1966).

5. O’Keefe, J. & Conway, D. H. Hippocampal place units in the freely moving rat: Why they fire where they fire. Exp. Brain Res. 31, 573–590 (1978).

6. Olton, D. S., Walker, J. A. & Gage, F. H. Hippocampal connections and spatial discrimination. Brain Res. 139, 295–308 (1978).

7. O’Keefe, J. & Dostrovsky, J. The hippocampus as a spatial map. Preliminary evidence from unit activity in the freely-moving rat. Brain Res. 34, 171–175 (1971).

8. Hafting, T., Fyhn, M., Molden, S., Moser, M.-B. & Moser, E. I. Microstructure of a spatial map in the entorhinal cortex. Nature 436, 801–806 (2005).

9. Muller, R. U. & Kubie, J. L. The effects of changes in the environment on the spatial firing of hippocampal complex-spike cells. J. Neurosci. Off. J. Soc. Neurosci. 7, 1951–1968 (1987).

10. Fyhn, M., Hafting, T., Treves, A., Moser, M.-B. & Moser, E. I. Hippocampal remapping and grid realignment in entorhinal cortex. Nature 446, 190–194 (2007).

11. O’Keefe, J. & Nadel, L. The Hippocampus as a Cognitive Map. Oxford University Press: Oxford, UK. (1978) (Oxford University Press, 1978).

12. Buzsáki, G. & Moser, E. I. Memory, navigation and theta rhythm in the hippocampal-entorhinal system. Nat. Neurosci. 16, 130–138 (2013).

13. Kacelnik, A., Brunner, D. & Gibbon, J. Timing Mechanisms in Optimal Foraging: Some Applications of Scalar Expectancy Theory. in Behavioural Mechanisms of Food Selection (ed. Hughes, R. N.) 61–82 (Springer Berlin Heidelberg, 1990). doi:10.1007/978-3-642-75118-9_4.

14. Marshall, A. T. & Kirkpatrick, K. Reinforcement learning models of risky choice and the promotion of risk-taking by losses disguised as wins in rats. J. Exp. Psychol. Anim. Learn. Cogn. 43, 262–279 (2017).

15. Brunner, D., Fairhurst, S., Stolovitzky, G. & Gibbon, J. Mnemonics for variability: remembering food delay. J. Exp. Psychol. Anim. Behav. Process. 23, 68–83 (1997).

16. Issa, J. B., Tocker, G., Hasselmo, M. E., Heys, J. G. & Dombeck, D. A. Navigating Through Time: A Spatial Navigation Perspective on How the Brain May Encode Time. Annu. Rev. Neurosci. 43, 73–93 (2020).

17. Heys, J. G., Wu, Z., Allegra Mascaro, A. L. & Dombeck, D. A. Inactivation of the Medial Entorhinal Cortex Selectively Disrupts Learning of Interval Timing. Cell Rep. 32, 108163 (2020).

18. Vo, A. et al. Medial entorhinal cortex lesions produce delay-dependent disruptions in memory for elapsed time. Neurobiol. Learn. Mem. 185, 107507 (2021).

19. Dias, M., Ferreira, R. & Remondes, M. Medial Entorhinal Cortex Excitatory Neurons Are Necessary for Accurate Timing. J. Neurosci. 41, 9932–9943 (2021).

20. Heys, J. G. & Dombeck, D. A. Evidence for a subcircuit in medial entorhinal cortex representing elapsed time during immobility. Nat. Neurosci. 21, 1574–1582 (2018).

21. Morris, R. G., Garrud, P., Rawlins, J. N. & O’Keefe, J. Place navigation impaired in rats with hippocampal lesions. Nature 297, 681–683 (1982).

22. Mishkin, M. Memory in monkeys severely impaired by combined but not by separate removal of amygdala and hippocampus. Nature 273, 297–298 (1978).

23. Squire, L. R. & Zola-Morgan, S. The medial temporal lobe memory system. Science 253, 1380–1386 (1991).

24. Otto, T. & Eichenbaum, H. Neuronal activity in the hippocampus during delayed non-match to sample performance in rats: evidence for hippocampal processing in recognition memory. Hippocampus 2, 323–334 (1992).

25. Verhagen, J. V., Wesson, D. W., Netoff, T. I., White, J. A. & Wachowiak, M. Sniffing controls an adaptive filter of sensory input to the olfactory bulb. Nat. Neurosci. 10, 631–639 (2007).

26. Heys, J. G., Rangarajan, K. V. & Dombeck, D. A. The functional micro-organization of grid cells revealed by cellular-resolution imaging. Neuron 84, 1079–1090 (2014).

27. Kraus, B. J. et al. During Running in Place, Grid Cells Integrate Elapsed Time and Distance Run. Neuron 88, 578–589 (2015).

28. Campbell, M. G., Attinger, A., Ocko, S. A., Ganguli, S. & Giocomo, L. M. Distance-tuned neurons drive specialized path integration calculations in medial entorhinal cortex. Cell Rep. 36, 109669 (2021).

29. Turi, G. F. et al. Vasoactive Intestinal Polypeptide-Expressing Interneurons in the Hippocampus Support Goal-Oriented Spatial Learning. Neuron 101, 1150–1165.e8 (2019).

30. Hardcastle, K., Maheswaranathan, N., Ganguli, S. & Giocomo, L. M. A Multiplexed, Heterogeneous, and Adaptive Code for Navigation in Medial Entorhinal Cortex. Neuron 94, 375–387.e7 (2017).

31. McDonald, R. J. & White, N. M. A triple dissociation of memory systems: hippocampus, amygdala, and dorsal striatum. Behav. Neurosci. 107, 3–22 (1993).

32. Toda, K. et al. Nigrotectal Stimulation Stops Interval Timing in Mice. Curr. Biol. CB 27, 3763–3770.e3 (2017).

33. Burak, Y. & Fiete, I. R. Accurate path integration in continuous attractor network models of grid cells. PLoS Comput. Biol. 5, e1000291 (2009).

34. Yoon, K. et al. Specific evidence of low-dimensional continuous attractor dynamics in grid cells. Nat. Neurosci. 16, 1077–1084 (2013).

35. Stensola, H. et al. The entorhinal grid map is discretized. Nature 492, 72–78 (2012).

36. Gardner, R. J. et al. Toroidal topology of population activity in grid cells. Nature 602, 123– 128 (2022).

37. Matell, M. S., Meck, W. H. & Nicolelis, M. A. L. Interval timing and the encoding of signal duration by ensembles of cortical and striatal neurons. Behav. Neurosci. 117, 760–773 (2003).

38. Hinton, S. C. & Meck, W. H. Frontal-striatal circuitry activated by human peak-interval timing in the supra-seconds range. Brain Res. Cogn. Brain Res. 21, 171–182 (2004).

39. Meck, W. H. Neuroanatomical localization of an internal clock: a functional link between mesolimbic, nigrostriatal, and mesocortical dopaminergic systems. Brain Res. 1109, 93–107 (2006).

40. Jin, D. Z., Fujii, N. & Graybiel, A. M. Neural representation of time in cortico-basal ganglia circuits. Proc. Natl. Acad. Sci. U. S. A. 106, 19156–19161 (2009).

41. Jazayeri, M. & Shadlen, M. N. A Neural Mechanism for Sensing and Reproducing a Time Interval. Curr. Biol. CB 25, 2599–2609 (2015).

42. Mello, G. B. M., Soares, S. & Paton, J. J. A scalable population code for time in the striatum. Curr. Biol. CB 25, 1113–1122 (2015).

43. Bakhurin, K. I. et al. Differential Encoding of Time by Prefrontal and Striatal Network Dynamics. J. Neurosci. 37, 854–870 (2017).

44. Meck, W. H., Church, R. M. & Olton, D. S. Hippocampus, time, and memory. Behav. Neurosci. 98, 3–22 (1984).

45. Jacobs, N. S., Allen, T. A., Nguyen, N. & Fortin, N. J. Critical role of the hippocampus in memory for elapsed time. J. Neurosci. Off. J. Soc. Neurosci. 33, 13888–13893 (2013).

46. Fortin, N. J., Agster, K. L. & Eichenbaum, H. B. Critical role of the hippocampus in memory for sequences of events. Nat. Neurosci. 5, 458–462 (2002).

47. Allen, T. A., Salz, D. M., McKenzie, S. & Fortin, N. J. Nonspatial Sequence Coding in CA1 Neurons. J. Neurosci. Off. J. Soc. Neurosci. 36, 1547–1563 (2016).

48. MacDonald, C. J., Lepage, K. Q., Eden, U. T. & Eichenbaum, H. Hippocampal ‘time cells’ bridge the gap in memory for discontiguous events. Neuron 71, 737–749 (2011).

49. Pastalkova, E., Itskov, V., Amarasingham, A. & Buzsáki, G. Internally generated cell assembly sequences in the rat hippocampus. Science 321, 1322–1327 (2008).

50. Manns, J. R., Howard, M. W. & Eichenbaum, H. Gradual changes in hippocampal activity support remembering the order of events. Neuron 56, 530–540 (2007).

51. Mankin, E. A. et al. Neuronal code for extended time in the hippocampus. Proc. Natl. Acad. Sci. U. S. A. 109, 19462–19467 (2012).

52. Tsao, A. et al. Integrating time from experience in the lateral entorhinal cortex. Nature 561, 57–62 (2018).

53. Issa, J. B. & Zhang, K. Universal conditions for exact path integration in neural systems. Proc. Natl. Acad. Sci. U. S. A. 109, 6716–6720 (2012).

54. Pachitariu, M. et al. Suite2p: beyond 10,000 neurons with standard two-photon microscopy. 061507 Preprint at 10.1101/061507 (2017).

55. Dombeck, D. A., Harvey, C. D., Tian, L., Looger, L. L. & Tank, D. W. Functional imaging of hippocampal place cells at cellular resolution during virtual navigation. Nat. Neurosci. 13, 1433–1440 (2010).

56. Skaggs, W., McNaughton, B. & Gothard, K. An Information-Theoretic Approach to Deciphering the Hippocampal Code. in Advances in Neural Information Processing Systems vol. 5 (Morgan-Kaufmann, 1992).

57. Climer, J. R. & Dombeck, D. A. Information Theoretic Approaches to Deciphering the Neural Code with Functional Fluorescence Imaging. eNeuro 8, ENEURO.0266-21.2021 (2021).

58. Park, E. H., Keeley, S., Savin, C., Ranck, J. B. & Fenton, A. A. How the Internally Organized Direction Sense Is Used to Navigate. Neuron 101, 285–293.e5 (2019).

